# *Drosophila* SUMM4 complex couples insulator function and DNA replication timing control

**DOI:** 10.1101/2021.10.02.462895

**Authors:** Evgeniya N. Andreyeva, Alexander V. Emelyanov, Markus Nevil, Lu Sun, Elena Vershilova, Christina A. Hill, Michael-C. Keogh, Robert J. Duronio, Arthur I. Skoultchi, Dmitry V. Fyodorov

## Abstract

Asynchronous replication of chromosome domains during S phase is essential for eukaryotic genome function, but the mechanisms establishing which domains replicate early versus late in different cell types remain incompletely understood. *Drosophila* SNF2-related factor SUUR imparts under- replication of late-replicating intercalary heterochromatin in polytene chromosomes. SUUR negatively regulates DNA replication fork progression; however, its mechanism of action remains obscure. Here we developed a novel method termed MS-Enabled Rapid protein Complex Identification (MERCI) to isolate a stable stoichiometric native complex SUMM4 that comprises SUUR and a chromatin boundary protein Mod(Mdg4)-67.2. Mod(Mdg4) stimulates SUUR ATPase activity and is required for a normal spatiotemporal distribution of SUUR *in vivo*. SUUR and Mod(Mdg4)-67.2 together mediate the activities of *gypsy* insulator that prevent certain enhancer-promoter interactions and establish euchromatin-heterochromatin barriers in the genome. Furthermore, *SuUR* or *mod(mdg4)* mutations reverse under-replication of intercalary heterochromatin. Thus, SUMM4 can impart late replication of intercalary heterochromatin by attenuating the progression of replication forks through euchromatin/heterochromatin boundaries. Our findings reveal that DNA replication can be delayed by a chromatin barrier and uncover a critical role for architectural proteins in replication control. They suggest a mechanism for replication timing that does not depend on an asynchronous firing of replication origins.

## Introduction

Replication of metazoan genomes occurs according to a highly coordinated spatiotemporal program, where discrete chromosomal regions replicate at distinct times during S phase (Rhind & Gilbert, 2013). The replication program follows the spatial organization of the genome in Megabase-long constant timing regions interspersed by timing transition regions (Marchal, Sima, & Gilbert, 2019). The spatiotemporal replication program exhibits correlations with genetic activity, epigenetic marks and features of 3D genome architecture and sub-nuclear localization. Yet the reasons for these correlations remain obscure. Interestingly, the timing of firing for any individual origin of replication is established during G1 before pre-replicative complexes (pre-RC) are assembled at origins (Dimitrova & Gilbert, 1999), suggesting a mechanism that involves factors other than the core replication machinery.

Most larval tissues of *Drosophila melanogaster* grow via G-S endoreplication cycles that duplicate DNA without cell division resulting in polyploidy (Zielke, Edgar, & DePamphilis, 2013). Endoreplicated DNA molecules frequently align in register to form giant polytene chromosomes (Zhimulev et al., 2004). Importantly, in some cell types, genomic domains corresponding to the latest replicated regions of dividing cells, specifically pericentric (PH) and intercalary (IH) heterochromatin, fail to fully replicate during each endocycle resulting in underreplication (UR). In both dividing and endoreplicating cells, these regions are depleted of sites for binding the Origin of Replication Complex (ORC) and thus their replication primarily relies on forks progressing from external origins (Sher et al., 2012). Although cell cycle programs are dissimilar between endoreplicating and mitotically dividing cells, they share biochemically identical DNA replication machinery (Zielke et al., 2013).

Thus, UR provides a facile readout for late replication initiation and delayed fork progression. The *Suppressor of Under-Replication* (*SuUR*) gene is essential for polytene chromosome UR in IH and PH (Belyaeva et al., 1998). In *SuUR* mutants, the DNA copy number in UR regions is equivalent to fully polyploidized regions of the genome. *SuUR* encodes a protein (SUUR) containing a helicase domain with homology to that of the SNF2/SWI2 family. The occupancy of ORC in IH and PH is not increased in *SuUR* mutants (Sher et al., 2012), and thus the increased replication of UR regions is likely not due to the firing of additional origins. Rather, SUUR negatively regulates the rate of replication fork progression (Nordman et al., 2014) by an unknown mechanism. It has been proposed (Posukh, Maksimov, Skvortsova, Koryakov, & Belyakin, 2015) that retardation of the replisome by SUUR takes place via simultaneous physical association with the components of the fork (*e.g*., CDC45 and PCNA) (Kolesnikova et al., 2013; Nordman et al., 2014) and repressive chromatin proteins, such as HP1a (Pindyurin et al., 2008).

Using a newly developed proteomics approach, we discovered that SUUR forms a stable complex with a chromatin boundary protein Mod(Mdg4)-67.2. We demonstrate that SUUR and Mod(Mdg4)-67.2 together are required for maximal UR of IH and full activity of the *gypsy* insulator, thereby implicating insulators in obstructing replisome progression and the control of late DNA replication.

## Results

### Identification of SUMM4, the native form of SUUR in *Drosophila* embryos

To determine how SUUR functions in replication control we sought to identify its native complex. Previous attempts to characterize the native form of SUUR by co-IP or tag-affinity purification gave rise to multiple putative binding partners (Kolesnikova et al., 2013; Munden et al., 2018; Nordman et al., 2014; Pindyurin et al., 2008). However, evaluating whether any of these proteins are present in a native SUUR complex is problematic because of the low abundance of SUUR, which also precludes its purification by conventional chromatography. Therefore, we developed a novel biochemical approach using embryonic extracts (which can be obtained in large quantities) that relies on partial purification by multi-step FPLC and shotgun proteomics of chromatographic fractions by quantitative LCMS (***Figure 1A***). We term this technology MERCI for MS-Enabled Rapid protein Complex Identification (*Materials and Methods*).

**Figure 1.**
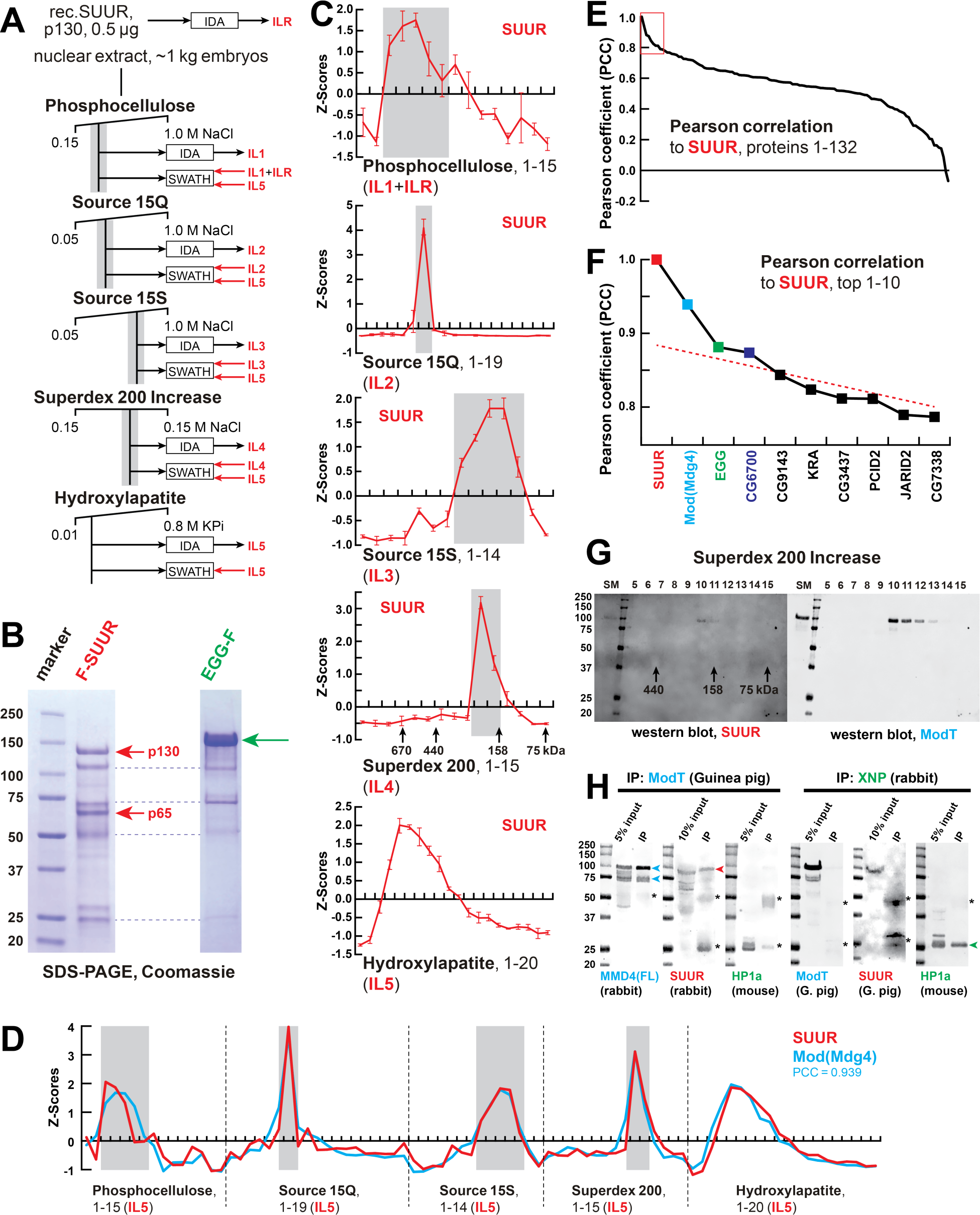
Identification of the SUMM4 complex by MERCI. (A) Schematic of FPLC purification of the native form of SUUR using MERCI approach. ILR, ion library obtained by IDA of recombinant FLAG- SUUR; IL1-5, ion libraries obtained by IDA of FPLC fractions from chromatographic steps 1-5. KPi, potassium phosphate, pH 7.6. (B) Recombinant FLAG-SUUR expressed in Sf9 cells. Identities of eight most prominent bands were determined by mass-spectroscopy. p130 and p65 correspond to full-length and C-terminally truncated FLAG-SUUR, respectively (red arrows). Other bands represent common Sf9-specific contaminants purified by FLAG chromatography (blue dashed lines), *cf* purified EGG-F (green arrow). Molecular mass marker bands are indicated (kDa). (C) SWATH quantitation profiles of SUUR fractionation across individual FPLC steps. Ion libraries (IL) used for SWATH quantitation are shown at the bottom. Z-scores across indicated column fractions are plotted; error bars, standard deviations (*N*=3). Gray rectangles, fraction ranges used for the next FPLC step; in Superdex 200 step, black arrows, expected peaks of globular proteins with indicated molecular masses in kDa. (D) SWATH quantitation profiles of SUUR (red) and Mod(Mdg4) (cyan) fractionation across five FPLC steps. IL5 ion library was used for SWATH quantification. (E) Pearson correlation of fractionation profiles for individual 132 proteins to that of SUUR, sorted from largest to smallest. Red box, the graph portion shown in (F). (F) Top ten candidate proteins with the highest Pearson correlation to SUUR. Red dashed line, trend line extrapolated by polynomial regression (*n* = 5) from the bottom 130 proteins. (G) Western blot analyses of Superdex 200 fractions with SUUR and ModT antibodies. Molecular mass markers are shown on the left (kDa). (H) Co-IP experiments. SUUR (red arrowhead) co-purifies from nuclear extracts with Mod(Mdg4)-67.2 (cyan arrowheads) but not HP1a (green arrowhead). Anti-XNP co-IPs HP1a but not SUUR of Mod(Mdg4)-67.2. Asterisks, IgG heavy and light chains detected due to antibody cross-reactivity. Mod(Mdg4)-67.2(FL) antibody recognizes all splice forms of Mod(Mdg4). Figure supplement 1. Identification of the SUMM4 complex by MERCI. Figure supplement 2. Quantification of SUUR in chromatographic fractions. Figure 1−table 1. UR domains and UR suppression in SUMM4 subunit mutant alleles.

The depth of proteomic quantification is limited by the range of peptides identified in the information-dependent acquisition (IDA), dubbed “ion library” (IL). Unfortunately, SUUR-specific peptides could not be found in ILs obtained from acquisitions of crude nuclear extracts or fractions from the first, phosphocellulose, step (IL1, ***Figure 1-figure supplement 1A****, **Data S1***). Thus, to quantify SUUR in phosphocellulose fractions, we augmented IL1 with the IL obtained by acquisition of recombinant SUUR (ILR, ***Figure 1A&B***). In ILs from subsequent chromatographic steps, peptides derived from native SUUR were detected (***Figure 1-figure supplement 1A****, **Data S1***) and used for quantification of cognate data-independent acquisitions (DIA/SWATH) (***Figure 1C***).

The final aspect of the MERCI algorithm calls for re-quantification of FPLC fraction SWATH acquisitions with an IL from the last step (IL5) that is enriched for peptides derived from SUUR and co-purifying polypeptides (***Figure 1A***) and includes only 140 proteins (***Figure 1-figure supplement 1A****, **Data S1***). In this fashion, scarce polypeptides (including SUUR and, potentially, SUUR-binding partners) that may not be detectable in earlier steps will not evade quantification. Purification profiles of proteins quantified in all five FPLC steps (132) were then artificially stitched into 83-point arrays of Z-scores (***Figure 1D, Data S2***). These profiles were Pearson-correlated with that of SUUR and ranked down from the highest Pearson coefficient, PCC (***Figure 1E***). Whereas the PCC numbers for the bottom 130 proteins lay on a smooth curve, the top two proteins, SUUR (PCC = 1.000) and Mod(Mdg4) (PCC = 0.939) fell above the extrapolated (by polynomial regression) curve (***Figure 1F***). Consistently, SUUR and Mod(Mdg4) exhibited nearly identical purification profiles in all five FPLC steps (***Figure 1D***), unlike the next two top-scoring proteins, EGG (PCC = 0.881) and CG6700 (PCC = 0.874) (***Figure 1-figure supplement 1B&C***). Also, HP1a (PCC = 0.503), which had been proposed to form a complex with SUUR (Pindyurin et al., 2008) did not co-purify with SUUR in any FPLC steps (***Figure 1-figure supplement 1D***).

Mod(Mdg4) is a BTB/POZ domain protein that functions as an adaptor for architectural proteins that promote various aspects of genome organization (Georgiev & Gerasimova, 1989; Gerasimova, Gdula, Gerasimov, Simonova, & Corces, 1995). It is expressed as 26 distinct polypeptides generated by splicing *in trans* of a common 5’-end precursor RNA with 26 unique 3’-end precursors (Buchner et al., 2000). IL5 contained seven peptides derived from Mod(Mdg4) (99% confidence). Whereas four of them mapped to the common N-terminal 402 residues, three were specific to the C-terminus of a particular form, Mod(Mdg4)-67.2 (***Figure 1-figure supplement 1E-G***). Peptides specific to other splice forms were not detected. We raised an antibody to the C-terminus of Mod(Mdg4)-67.2, designated ModT antibody, and analyzed size exclusion column fractions by immunoblotting.

Consistent with SWATH analyses (***Figure 1C&D***) SUUR and Mod(Mdg4)-67.2 polypeptides copurified as a complex with an apparent molecular mass of ∼250 kDa (***Figure 1G***). Finally, we confirmed that SUUR specifically co-immunoprecipitated with Mod(Mdg4)-67.2 from embryonic nuclear extracts (***Figure 1H***). As a control, XNP co-immunoprecipitated with HP1a as shown previously (Emelyanov, Konev, Vershilova, & Fyodorov, 2010), but did not with SUUR or Mod(Mdg4) (***Figure 1H***). We conclude that SUUR and Mod(Mdg4) form a stable stoichiometric complex that we term SUMM4.

### Biochemical activities of recombinant SUMM4 *in vitro*

We reconstituted recombinant SUMM4 complex by co-expressing FLAG-SUUR with Mod(Mdg4)-67.2-His_6_ in Sf9 cells and purified it by FLAG affinity chromatography (***Figure 2A***). Mod(Mdg4)-67.2 is the predominant form of Mod(Mdg4) expressed in embryos (*e.g*., ***Figure 1H***). Thus, minor Mod(Mdg4) forms may have failed to be identified by IDA in IL5 (***Figure 1-figure supplement 1E***). We discovered that FLAG-SUUR did not co-purify with another splice form, Mod(Mdg4)-59.1 (***Figure 1-figure supplement 1G***, ***Figure 2A***). Whereas the identity of an ∼100- kDa Mod(Mdg4)-67.2-His_6_ band copurifying with FLAG-SUUR was confirmed by mass-spec sequencing, the FLAG-purified material from Sf9 cells expressing FLAG-SUUR and Mod(Mdg4)-59.1 did not contain Mod(Mdg4)-specific peptides. Therefore, the shared N-terminus of Mod(Mdg4) (1-402) is not sufficient for interactions with SUUR. However, this result does not exclude a possibility that SUUR may form complex(es) with some of the other, low-abundance 24 splice forms of Mod(Mdg4). The SUUR- Mod(Mdg4)-67.2 interaction is specific, as the second-best candidate from our correlation analyses (*Drosophila* SetDB1 ortholog EGG; ***Figure 1F***) did not form a complex with FLAG-SUUR (***Figure 2-figure supplement 1A***), although it associated with its known partner WDE, an ortholog of hATF7IP/mAM (Wang et al., 2003).

**Figure 2.**
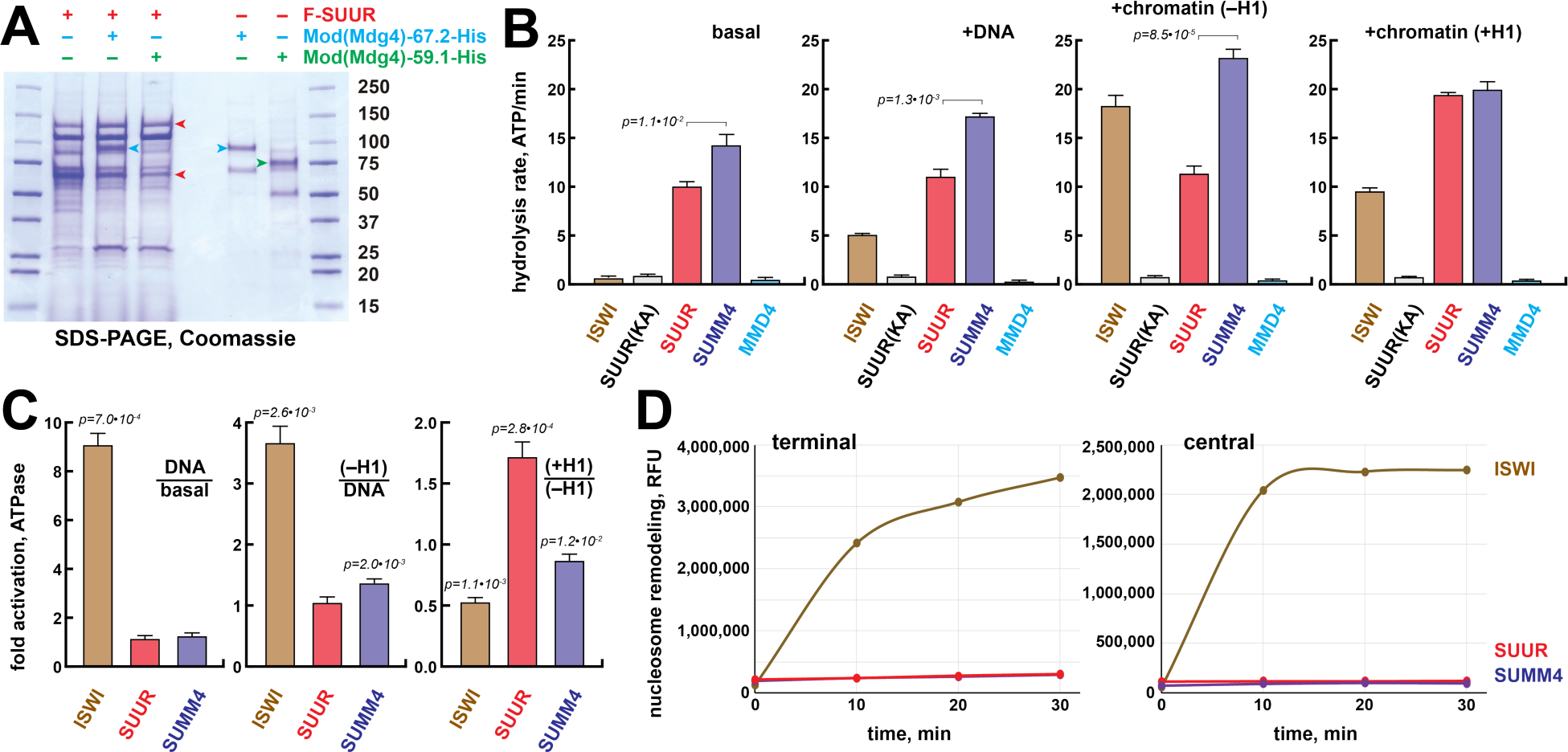
Biochemical activities of SUMM4. (A) Recombinant SUMM4. Mod(Mdg4)-His_6_, 67.2 (p100, cyan arrowhead) and 59.1 (p75, green arrowhead) splice forms were co-expressed with FLAG-SUUR (red arrowheads, p130 and p65) or separately in Sf9 cells and purified by FLAG or Ni-NTA affinity chromatography. Mod(Mdg4)-67.2 forms a specific complex with SUUR. Identities of the 130-, 100-, 75- and 65-kDa protein bands from FLAG- and Ni-NTA-purified material were determined by mass-spectroscopy. (B) ATPase activities of recombinant ISWI (brown bars), FLAG-SUUR (red bars) and SUMM4 (FLAG-SUUR + Mod(Mdg4)-67.2-His_6_, purple bars). Equimolar amounts of proteins were analyzed in reactions in the absence or presence of plasmid DNA or equivalent amounts of reconstituted oligonucleosomes, ±H1. SUUR(KA) and MMD4, ATPases activities of K59A mutant of SUUR (gray bars) and Mod(Mdg4)-67.2-His_6_ (cyan bars). Hydrolysis rates were converted to moles ATP per mole protein per minute. All reactions were performed in triplicate, error bars represent standard deviations. *p*-values for statistically significant differences are indicated (Mann-Whitney test). (C) DNA- and nucleosome-dependent stimulation or inhibition of ATPase activity. The activities were analyzed as in (B). Statistically significant differences are shown (Mann-Whitney test). (D) Nucleosome sliding activities by EpiDyne^®^-PicoGreen™ assay (see *Materials and Methods*) with 5 nM of recombinant ISWI, SUUR or SUMM4. Reaction time courses are shown for terminally (6-N-66) and centrally (50-N-66) positioned mononucleosomes (Figure 2***-figure supplement 1G****-**J***). RFU, relative fluorescence units produced by PicoGreen fluorescence. Figure supplement 1. Recombinant proteins, substrates and biochemical assays.

The N-terminus of SUUR contains a region homologous with SNF2-like DEAD/H helicase domains. Although SUUR requires its N-terminal domain to function *in vivo* (Munden et al., 2018), it has been hypothesized to be inactive as an ATPase (Nordman & Orr-Weaver, 2015). We analyzed the ability of recombinant SUUR and SUMM4 (***Figure 2A***) to hydrolyze ATP *in vitro* in comparison to recombinant *Drosophila* ISWI (***Figure 2-figure supplement 1B***). Purified recombinant Mod(Mdg4)-67.2 and a variant SUUR protein with a point mutation in the putative Walker A motif (K59A) were used as negative controls (***Figure 2A***, ***Figure 2-figure supplement 1B***). Contrary to the prediction, both SUUR and SUMM4 exhibited strong ATPase activities (***Figure 2B***). SUMM4 was 1.4- to 2-fold more active than SUUR alone, indicating that Mod(Mdg4)-67.2 stimulates SUUR enzymatic activity. We then examined whether DNA and nucleosomes can stimulate the activity of SUUR. To this end, we reconstituted oligonucleosomes on plasmid DNA (***Figure 2-figure supplement 1B-E***). Linker histone H1-containing chromatin was also used as a substrate/cofactor, because SUUR has been demonstrated to physically interact with H1 (Andreyeva et al., 2017). In contrast to ISWI, SUUR was not stimulated by addition of DNA or nucleosomes and moderately (by about 70%) activated by H1- containing oligonucleosomes (***Figure 2C***) consistent with its reported direct physical interaction with H1 (Andreyeva et al., 2017).

We examined the nucleosome remodeling activities of SUUR and SUMM4; specifically, their ability to expose a positioned DNA motif in the EpiDyne^®^-PicoGreen™ assay (*Materials and Methods* and ***Figure 2-figure supplement 1F***). Centrally or terminally positioned mononucleosomes were efficiently mobilized by ISWI and human BRG1 in a concentration- and time-dependent manner (***Figure 2-figure supplement 1G-J***). In contrast, SUUR and SUMM4 did not reposition either nucleosome (***Figure 2D***). Thus, SUUR and SUMM4 do not possess a detectable remodeling activity and may resemble certain other SNF2-like enzymes (*e.g*., RAD54) that utilize the energy of ATP hydrolysis to mediate alternate DNA translocation reactions (Jaskelioff, Van Komen, Krebs, Sung, & Peterson, 2003).

### The distribution of SUMM4 complex *in vivo*

We examined the positions of SUUR and Mod(Mdg4)-67.2 within polytene chromosomes by indirect immunofluorescence (IF) and discovered that they overlap at numerous locations (***Figure 3A***, ***Figure 3-figure supplement 1A&B***). In late endo-S phase when SUUR exhibited a characteristic distribution, it co-localized with Mod(Mdg4)-67.2 at numerous (hundreds of) loci along the chromosome arms (***Figure 3-figure supplement 1B***). Mod(Mdg4)-67.2 was present at classical regions of SUUR enrichment, such as UR domains in 75C and 89E (***Figure 3-figure supplement 1A***). The chromocenter, which consists of under-replicated PH, contains SUUR but did not show occupancy by Mod(Mdg4)-67.2 (***Figure 3-figure supplement 1B***). Conversely, there were multiple sites of Mod(Mdg4)-67.2 localization that were free of SUUR (***Figure 3-figure supplement 1B***).

**Figure 3.**
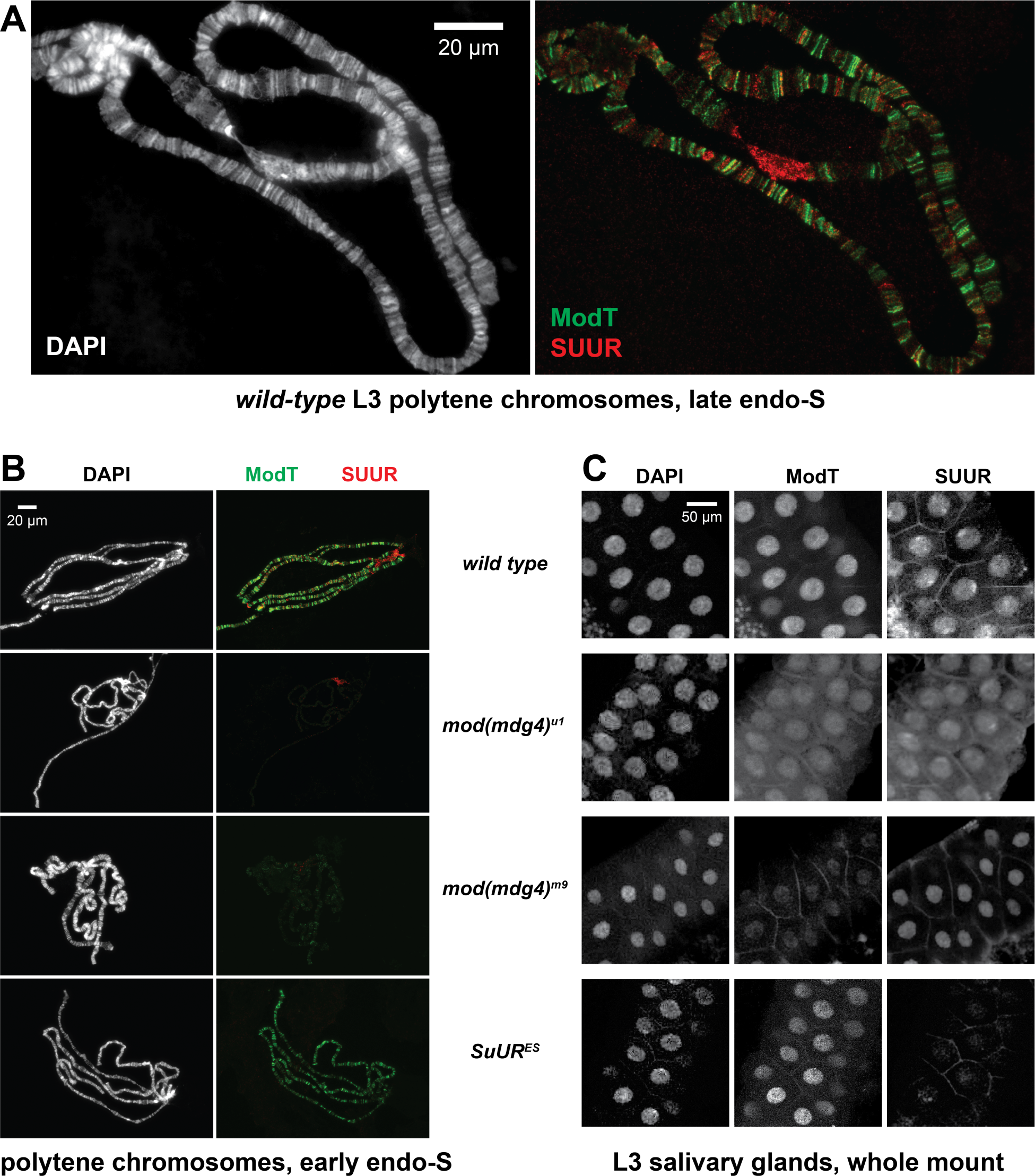
Spatiotemporal distribution of SUMM4 *in vivo*. (A) Colocalization of SUUR and Mod(Mdg4)-67.2 in *wild-type* polytene chromosomes. Localization patterns of Mod(Mdg4)-67.2 and SUUR in L3 polytene chromosomes were analyzed by indirect IF staining. The polytene spread fragment (3L and 3R arms) corresponds to a nucleus in late endo-S phase, according to PCNA staining (Figure 3***-figure supplement 1A***). ModT (green) and SUUR (red) signals overlap extensively in euchromatic arms. Note the additional strong ModT IF loci that are SUUR-free as well as Mod(Mdg4)- 67.2-free SUUR in pericentric 3LR. DAPI staining shows the overall chromosome morphology. (B) SUUR loading into chromosomes during early endo-S phase is compromised in *mod(mdg4)* mutants. *SuUR* mutation does not appreciably change the distribution of Mod(Mdg4)-67.2. Endo-S timing was established by PCNA staining (Figure 3***-figure supplement 1J***). (C) Abnormal subcellular distribution of SUMM4 subunits in *mod(mdg4)* and *SuUR* mutants. L3 salivary glands were fixed and whole-mount- stained with DAPI, ModT and SUUR antibodies. Whereas both polypeptides are mostly nuclear in wild type, they are partially mis-localized to the cytoplasm in *mod(mdg4)^u1^* mutant. Figure supplement 1. Spatiotemporal distribution of SUUR and Mod(Mdg4) *in vivo* and alternative complex(es) of Mod(Mdg4).

Individual pixel intensities of IF signals for SUUR and Mod(Mdg4)-67.2 were plotted as a 2D scatter plot (***Figure 3-figure supplement 1C***) and were found to exhibit a weak positive correlation (R^2^=0.278). Consistent with the possible multi-phasic relative distribution of SUUR and Mod(Mdg4)-67.2 (***Figure 3-figure supplement 1B***), the 2D plot encompassed four distinct areas, where SUUR and Mod(Mdg4)-67.2 were co-localized, enriched separately or absent (***Figure 3-figure supplement 1D***). When regions of SUUR-alone and Mod(mdg4)-67.2-alone enrichment were excluded, and only the regions of their apparent colocalization were considered, the anti-SUUR and anti-ModT signals exhibited a strong positive correlation (R^2^=0.568, ***Figure 3-figure supplement 1D***).

The existence of chromosome loci heavily enriched for Mod(Mdg4)-67.2 but devoid of SUUR suggests that there are additional native form(s) of Mod(Mdg4)-67.2, either as an individual polypeptide or in complex(es) other than SUMM4. When we fractionated *Drosophila* nuclear extract using a different progression of FPLC steps (***Figure 3-figure supplement 1E***), we found that Mod(Mdg4)-67.2 can form a megadalton-sized complex that did not contain SUUR (***Figure 3-figure supplement 1F-H***). Therefore, a more intricate pattern of Mod(Mdg4)-67.2 distribution likely reflects loading of both SUMM4 and an alternative Mod(Mdg4)-67.2-containing complex.

We tested whether SUUR and Mod(Mdg4) loading into polytene chromosomes were mutually dependent using mutant alleles of *SuUR* and *mod(mdg4)*. *SuUR^ES^* is a null allele of *SuUR* (Makunin et al., 2002). *mod(mdg4)^m9^* is a null allele with a deficiency that removes gene regions of the shared 5’- end precursor and eight specific 3’-precursors (Savitsky, Kim, Kravchuk, & Schwartz, 2016). *mod(mdg4)^u1^* contains an insertion of a *Stalker* element in the last coding exon of Mod(Mdg4)-67.2 3’- precursor (Gerasimova et al., 1995), and thus is predicted only to disrupt expression of this isoform.

*SuUR^ES^* and *mod(mdg4)^u1^* are homozygous viable, and *mod(mdg4)^m9^* is recessive adult pharate lethal. We could not detect Mod(Mdg4)-67.2 expression in homozygous *mod(mdg4)^m9^*L3 salivary glands by immunoblotting, whereas *mod(mdg4)^u1^* expressed a truncated polypeptide (*cf*, ∼70 kDa and ∼100 kDa, ***Figure 3-figure supplement 1I***). The truncated 70-kDa polypeptide failed to load into polytene chromosomes (***Figure 3B***, ***Figure 3-figure supplement 1J***). As shown previously, SUUR could not be detected in *SuUR^ES^*chromosomes. Since homozygous *mod(mdg4)^m9^* L3 larvae were produced by *inter se* crosses of heterozygous parents, the very low amounts of Mod(Mdg4)-67.2 in *mod(mdg4)^m9^* polytene chromosomes (barely above the detection limit) were presumably maternally contributed.

The absence (or drastic decrease) of Mod(Mdg4)-67.2 also strongly reduced the loading of SUUR (***Figure 3B***, ***Figure 3-figure supplement 1J***). The normal distribution pattern of SUUR in polytene chromosomes is highly dynamic (Andreyeva et al., 2017; Kolesnikova et al., 2013). SUUR is initially loaded in chromosomes at the onset of endo-S phase and then re-distributes through very late endo-S, when it accumulates in UR domains and PH. In both *mod(mdg4)* mutants, we observed a striking absence of SUUR in polytene chromosomes during early endo-S (***Figure 3B***), which indicates that the initial deposition of SUUR is dependent on its interactions with Mod(Mdg4). Although SUUR deposition slightly recovered by late endo-S, it was still several fold weaker than that in wild type.

Potentially, in the absence of Mod(Mdg4), SUUR may be tethered to IH and PH loci by direct binding with linker histone H1 as shown previously (Andreyeva et al., 2017). Finally, the gross subcellular distribution of SUUR also strongly correlated with that of Mod(Mdg4): a mis-localization of truncated Mod(Mdg4)-67.2 from nuclear to partially cytoplasmic was accompanied by a similar mis-localization of SUUR (***Figure 3C***). This result indicates that the truncation of Mod(Mdg4) in *mod(mdg4)^u1^*may have an antimorphic effect by mis-localization and deficient chromatin binding of interacting polypeptides, including SUUR (***Figure 3C***) and others (***Figure 3-figure supplement 1E****-**H***).

### The role of SUMM4 as an effector of the insulator/chromatin barrier function

Mod(Mdg4)-67.2 does not directly bind DNA but instead is tethered by a physical association with zinc finger factor Suppressor of Hairy Wing, Su(Hw). Su(Hw) directly binds to consensus sequences that are present in *gypsy* transposable elements and are also widely distributed across the *Drosophila* genome in thousands of copies (Adryan et al., 2007). Mod(Mdg4)-67.2 was previously shown to be essential for the insulator activity of *gypsy* (Gerasimova et al., 1995), which functions *in vivo* to prevent enhancer-promoter interactions and establish a barrier to the propagation of chromatin forms (Cai & Levine, 1995; Roseman, Pirrotta, & Geyer, 1993). We therefore tested whether SUMM4 contributes to *gypsy* insulator functions.

The *ct^6^* allele of *Drosophila* contains a *gypsy* element inserted between the wing enhancer and promoter of the gene *cut* that inactivates *cut* expression and results in abnormal wing development (***Figure 4A***). We discovered that both *mod(mdg4)^u1^* and *SuUR^ES^* mutations partially suppressed this phenotype (***Figure 4A***) and significantly increased the wing size compared to *ct^6^*allele alone (***Figure 4B***). Thus, both subunits of SUMM4 are required to mediate the full enhancer-blocking activity of *gypsy*. Another insulator assay makes use of a collection of *P{SUPor-P}* insertions that contain the *white* reporter flanked by 12 copies of *gypsy* Su(Hw)-binding sites. When *P{SUPor-P}* is inserted in heterochromatin, *white* is protected from silencing resulting in red eyes (Roseman et al., 1995). Both *mod(mdg4)^u1^* and *SuUR^ES^*relieved the chromatin barrier function of Su(Hw) sites, causing repression of *white* (***Figure 4C***). We conclude that SUMM4 is an insulator complex that contributes to the enhancer-blocking and chromatin boundary functions of *gypsy* by a mechanism schematized in ***Figure 4D&E***.

**Figure 4.**
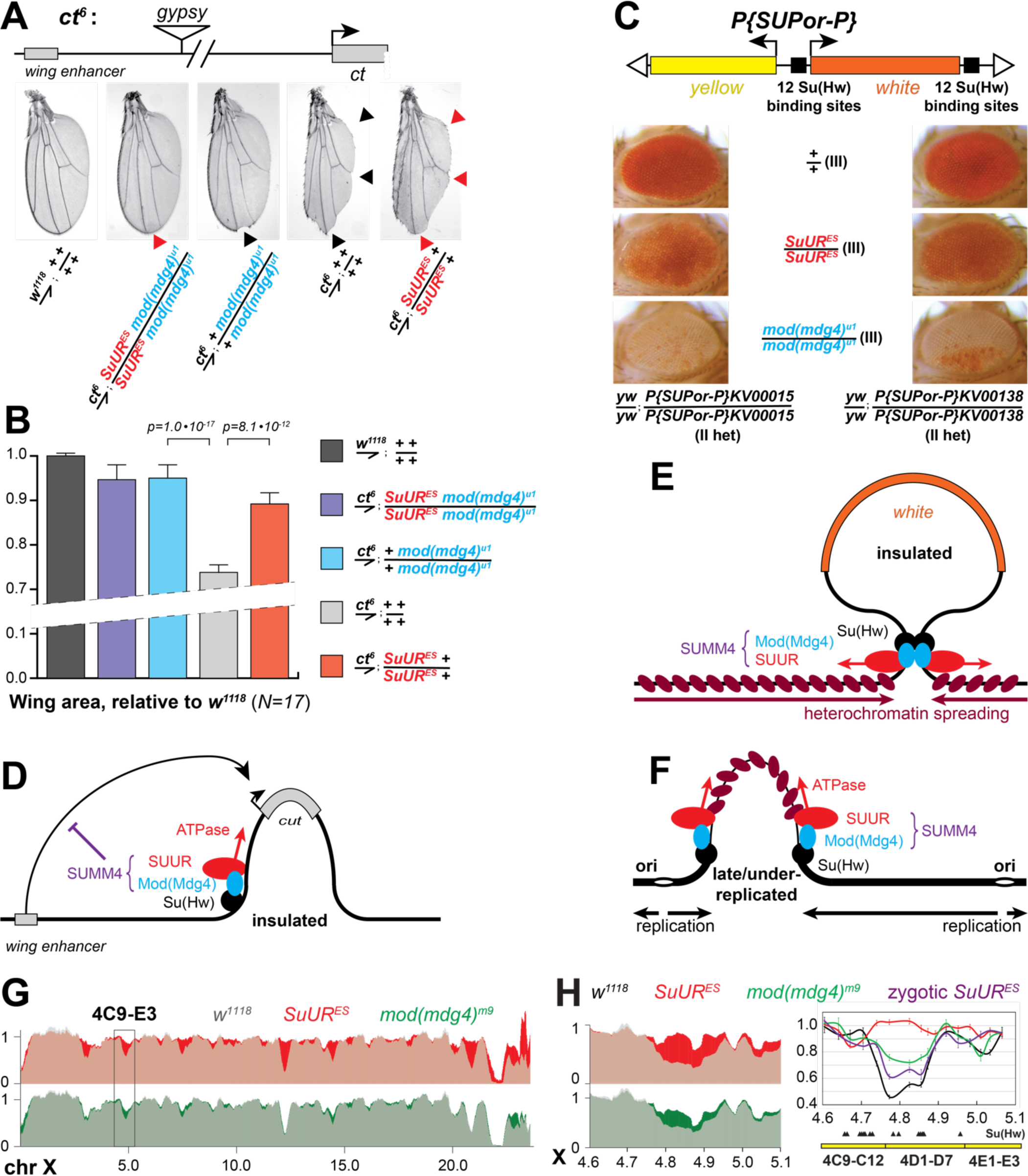
Biological functions of SUMM4 in regulation of gene expression and DNA replication. (A) SUMM4 subunits are required for the enhancer-blocking activity in *ct^6^*. Top: schematic diagram of the *ct^6^* reporter system; the *gypsy* retrotransposon is inserted in between the wing enhancer and promoter of *cut* (Bag, Dale, Palmer, & Lei, 2019). Bottom left: the appearance of wild type adult wing; bottom right: the appearance of *ct^6^* adult wing in the wild-type background. *SuUR^ES^*and *mod(mdg4)^u1^* alleles are recessive suppressors of the *ct^6^*phenotype. Red and black arrowheads point to distinct anatomical features of the wing upon *SuUR* mutation. (B) Relative sizes (areas) of wings in adult male flies of the indicated phenotypes were measured as described in *Materials and Methods*. *p*-values for statistically significant differences are indicated (t- test). (C) SUMM4 subunits are required for the chromatin barrier activity of Su(Hw) binding sites. Top: schematic diagram of the *P{SUPor-P}* reporter system (Bellen et al., 2004); clustered 12 copies of *gypsy* Su(Hw) binding sites flanks the transcription unit of *white*. *KV00015* and *KV00138* are *P{SUPor-P}* insertions in pericentric heterochromatin of 2L. *SuUR^ES^* and *mod(mdg4)^u1^* alleles are recessive suppressors of the boundary that insulates *white* from heterochromatin encroachment. (D) Schematic model for the function of SUMM4 in blocking enhancer-promoter interactions in the *ct^6^* locus. (E) Schematic model for the function of SUMM4 in establishing a chromatin barrier in heterochromatin-inserted *P{SUPor-P}* elements. (F) Schematic model for a putative function of SUMM4 in blocking/retardation of replication fork progression in IH domains. (G) Analyses of DNA copy numbers in *Drosophila* salivary gland cells. DNA from L3 salivary glands was subjected to high-throughput sequencing. DNA copy numbers (normalized to diploid embryonic DNA) are shown across the X chromosome. Genomic coordinates in Megabase pairs are indicated at the bottom. The control trace (*w^1118^* allele) is shown as semitransparent light gray in the foreground; *SuUR^ES^*(homozygous null) and *mod(mdg4)^m9^* (zygotic null from crosses of heterozygous parents) traces are shown in the background in red and green, respectively; their overlaps with *w^1118^*traces appear as lighter shades of colors. Black box, 4C9-E3 cytological region. (H) Close- up view of DNA copy numbers in region 4C9-E3 from high-throughput sequencing data are presented as in (G). DNA copy numbers were also measured independently by real-time qPCR. The numbers were calculated relative to embryonic DNA and normalized to a control intergenic region. The X-axis shows chromosome positions (in Megabase pairs) of target amplicons. Black, *w^1118^*; red, *SuUR^ES^* (homozygous null); green, *mod(mdg4)^m9^* (zygotic null from crosses of heterozygous parents); purple, *SuUR^ES^* (zygotic null from crosses of heterozygous parents). Error bars represent the confidence interval (see *Materials and Methods*). Black arrowheads, positions of mapped Su(Hw) binding sites (Negre et al., 2010). Yellow boxes show approximate boundaries of cytogenetic bands. Figure supplement 1. Biological functions of SUMM4 in regulation of under-replication. Figure 4−table 2. Primer sequences used for qPCR.

### The role of SUMM4 in regulation of DNA replication in polytene chromosomes

A similar, chromatin partitioning-related mechanism may direct the function of SUUR in the establishment of UR in late-replicating IH domains of polytene chromosomes (***Figure 4F***). It has been long known that 3D chromosome partitioning maps show an “uncanny alignment” with replication timing maps (Rhind & Gilbert, 2013). To examine the possible roles of SUMM4 in UR, we measured DNA copy number genome-wide in salivary glands of L3 larvae by next generation sequencing (NGS). In *w^1118^* control salivary glands, the DNA copy profile revealed large (>100-kbp) domains of reduced ploidy (***Figure 4-figure supplement 1A***), similar to a previous report (Andreyeva et al., 2017). Excluding pericentric and sub-telomeric heterochromatin, we called 70 UR regions (***Table 1***) in euchromatic arms, as described in *Materials and Methods*.

**Table 1.**
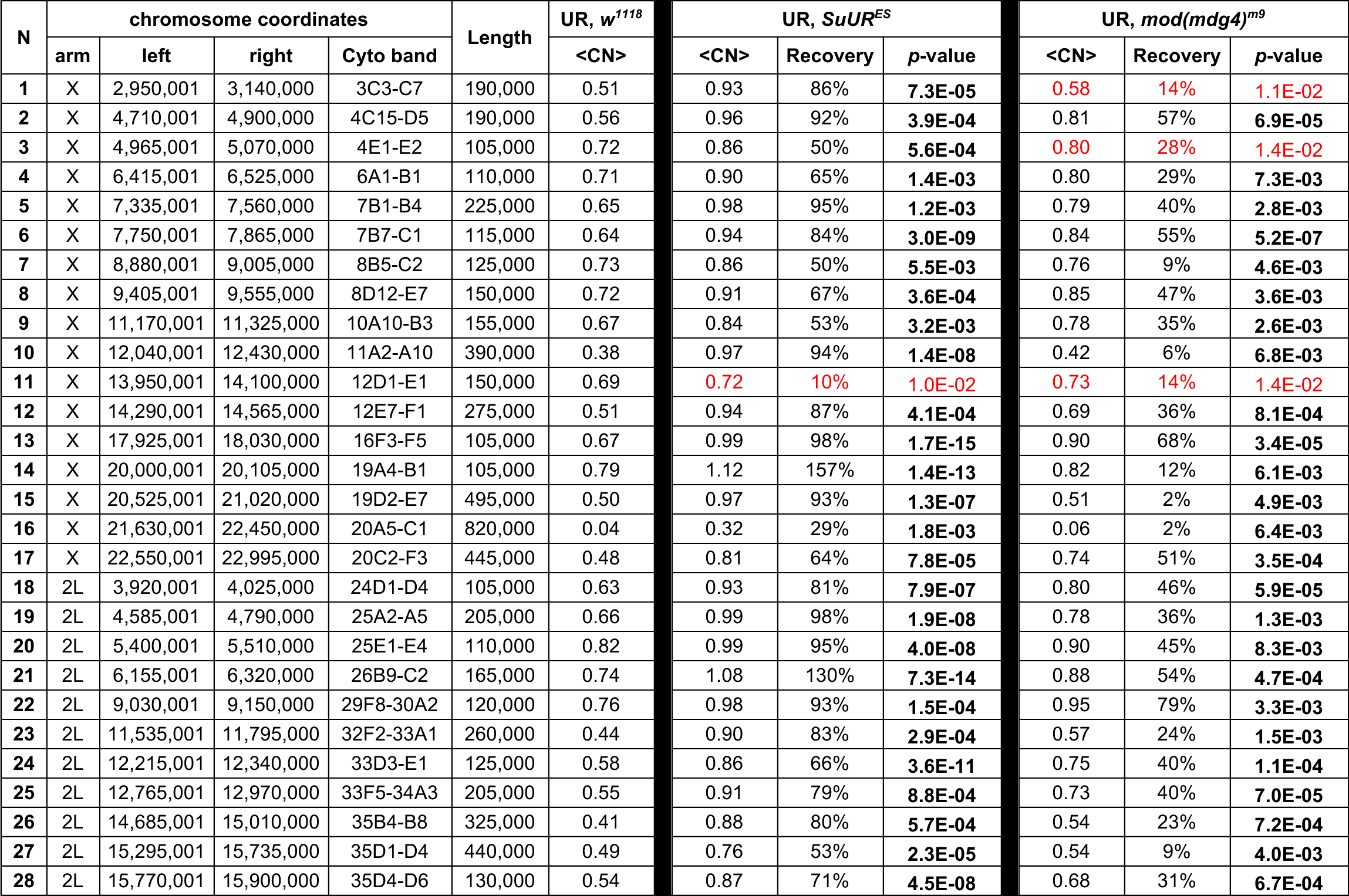
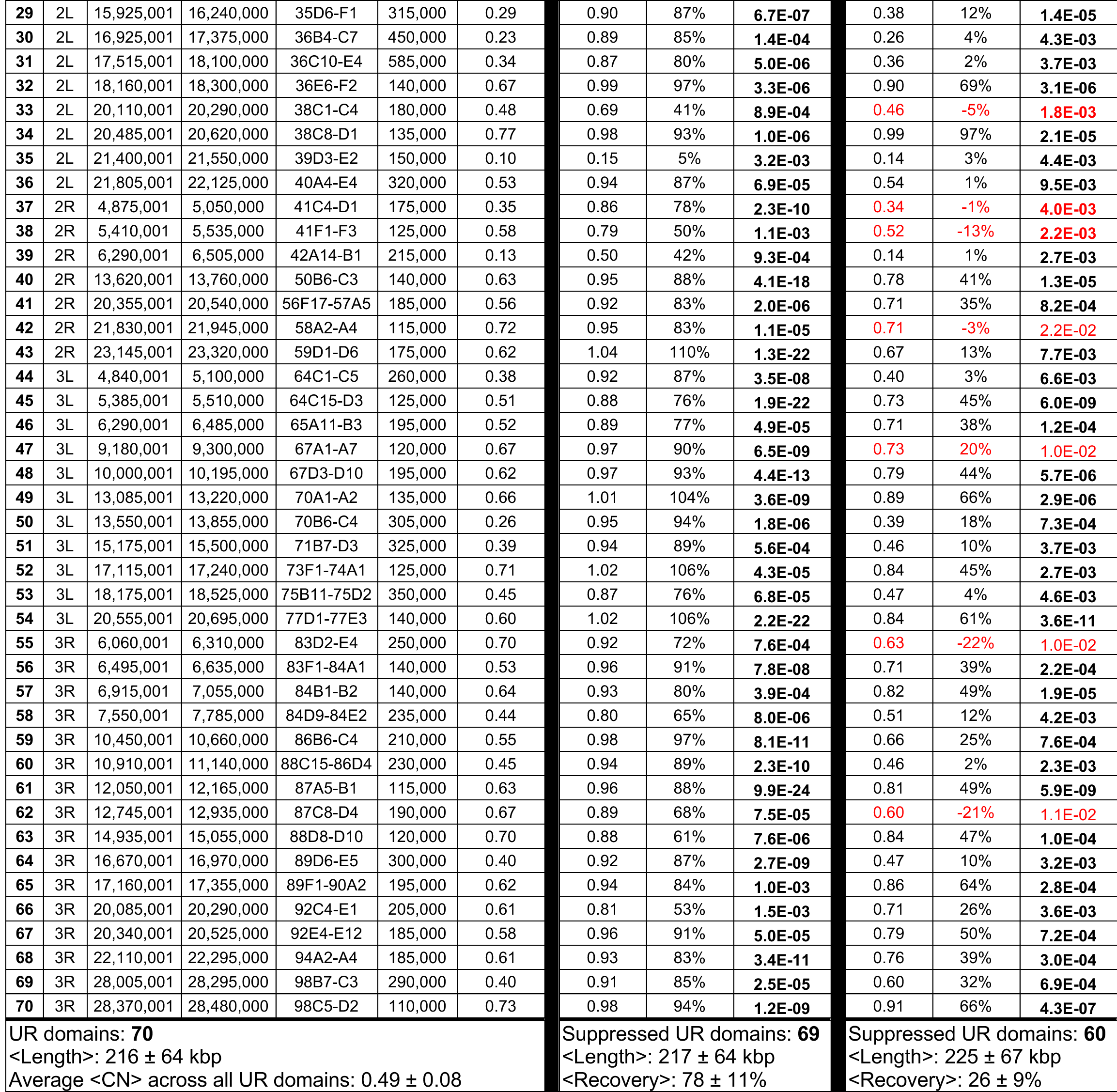
UR domains and UR suppression in SUMM4 subunit mutant alleles. Domains of underreplication (UR) in euchromatic arms of polytene chromosomes were called in *w^1118^*as described in *Methods*. Their genomic coordinates, approximate cytological location (“Cyto band”) and average DNA copy numbers (“<CN>”) in homozygous *w^1118^*, *SuUR^ES^* and *mod(mdg4)^m9^*L3 larvae are shown. <CN> numbers were normalized to the average DNA copy numbers across euchromatic genome. Percent UR recovery levels were calculated as (<CN>*mut* – <CN>*w1118*) / (1 – <CN>*w1118*); negative numbers indicate increased UR. UR *p*-values were calculated using the DESeq2 package by averaging the Wald test *p*-values of each 5-kbp bin significantly different than the *w^1118^* signal. UR was called as suppressible by a mutant if *p* < 0.01; regions that do not exhibit a statistically significant recovery of UR are marked in red. Averages of <CN> across all called UR domains and averages of percent Recovery across all suppressible UR domains (“<RECOVERY>”, bottom row) were adjusted for each UR domain length; calculation errors = standard deviations.

In both *SuUR* and *mod(mdg4)^m9^* null larvae, we observed statistically significant suppression of UR in IH (***Figure 4G***, ***Figure 4-figure supplement 1B***, ***Table 1***). Consistent with its lack of accumulation within the chromocenter of polytene chromosomes (***Figure 3A***), Mod(Mdg4) was dispensable for UR in PH. The NGS data strongly correlated with qPCR measurements of DNA copy numbers (***Figure 4H***, ***Figure 4-figure supplement 1C***). Furthermore, cytological evidence in the 75C region supported the molecular analyses in that both mutants exhibited a brighter DAPI staining of the 75C1-2 band than that in *w^1118^*, indicative of higher DNA content (***Figure 4-figure supplement 1C***).

Uniformly, *SuUR* mutation gave rise to a stronger relief of UR than that produced by the *mod(mdg4)^m9^* null allele (***Table 1***). This result can be explained by embryonic deposition of functional Mod(Mdg4) proteins and RNA by heterozygous mothers, unlike the complete absence of SUUR throughout the life cycle of the homozygous viable and fertile *SuUR^ES^*animals. Although third instar larvae are >1,000-fold larger, volume-wise, than the embryos, persistent Mod(Mdg4)-67.2 can still be detected in polytene chromosomes of these larvae by IF despite its dilution and degradation (***Figure 3C***, ***Figure 3-figure supplement 1J***). In contrast, unlike L3, first instar larvae are nearly identical in size to the embryos. Therefore, since the endoreplication cycles initiate in embryos and L1, in *mod(mdg4)^m9^* animals the first few out of 10-11 rounds of chromosome polytenization take place with an almost normal amount of Mod(Mdg4) present, which may substantially limit the effect of *mod(mdg4)^m9^* mutation on UR as measured in L3.

Seemingly, there is a contradiction between a strong effect that *mod(mdg4)* null mutation has on the loading of SUUR in polytene chromosomes (***Figure 3B***) and a weaker effect on UR (***Figure 4G&H*** and ***Figure 4-figure supplement 1B****-**D***). However, the SUUR occupancy is examined in L3 after maternal *mod(mdg4)* product is nearly eliminated (***Figure 3B***). On the other hand, the DNA copy number, although also measured in L3 (***Figure 4G&H*** and ***Figure 4-figure supplement 1B****-**D***), is a product of multiple rounds of endoreplication that initiate before Mod(Mdg4) is exhausted. To validate the putative effect of maternally contributed SUMM4 on the establishment of UR, we performed qPCR measurements of DNA copy numbers in salivary glands of homozygous *SuUR* animals produced by *inter se* crosses of heterozygous *SuUR^ES^/+* parents (***Figure 4H***, ***Figure 4-figure supplement 1C***, zygotic *SuUR^ES^*). Similar to the maternal Mod(Mdg4), the initial maternal contribution of SUUR partially limited the reversal of UR in cytological regions 4D and 75C. Thus, when the *SuUR* and *mod(mdg4)* null mutant animals are similarly derived from heterozygous mothers that deposit wild-type gene product into their progeny, the mutant UR phenotypes in the third instar larval salivary gland are essentially indistinguishable.

We conclude that SUUR and Mod(Mdg4)-67.2 act together as subunits of stable SUMM4 complex, which is required for the establishment of UR in the IH domains of *Drosophila* polytene chromosome.

## Discussion

### MERCI is a powerful new approach to characterize stable stoichiometric protein complexes

We present here a facile method, termed MERCI, to rapidly identify subunits of stable native complexes by only partial chromatographic purification. It allows one to circumvent the conventional, rate-limiting approach to purify proteins to apparent homogeneity. Since a multi-step FPLC scheme invariably leads to an exponential loss of material, reducing the number of purification steps in the MERCI protocol allows identification of rare complexes, such as SUMM4, which may be present in trace amounts in native sources. On the other hand, MERCI obviates introduction of false-positives frequently associated with tag purification of ectopically expressed targets that render results less reliable. Notably, MERCI is not limited to analyses of known polypeptides, since it is readily amenable to fractionation of native factors based on a correlation with their biochemical activities *in vitro*.

The dissection of protein interactome by extract fractionation on orthogonal FPLC columns and MS-based approaches has been previously attempted (Havugimana et al., 2012; Shatsky et al., 2016).

However, unlike the newly developed MERCI approach, these studies were aimed at comprehensive, proteome-wide analyses, which managed to only yield data for the most abundant complexes. The major distinction of the MERCI protocol is that it is targeted towards a particular protein (SUUR in this study). The crucial final stage of the MERCI algorithm is re-quantification of all acquired SWATH data using a library acquired from fractions of the last column (IL5, ***Figure 1A&D***). The target protein and co-purifying polypeptides are substantially enriched after several chromatographic steps and thus, yield a greater number of detected peptides, which helps a more precise quantification. Although SWATH allows reliable measurement of picogram amounts of proteins (***Figure 1-figure supplement 2A&B***), the range of quantified polypeptides is always limited by those present in IDA acquisitions (ion libraries). For low-abundance proteins, such as SUUR and Mod(Mdg4), specific peptides are not detectable by IDA in earlier chromatographic steps (***Data S1***). Consequently, SWATH quantification using only the cognate ion libraries would not discern the near perfect co-fractionation of SUUR and Mod(Mdg4) in all five steps (***Figure 1D***), precluding identification of the SUUR-Mod(Mdg4) complex (***Figure 1D****-**F***).

One limitation of the MERCI protocol is its failure to measure the absolute amounts of identified polypeptides. For instance, quantification of SWATH data (***Figure 1C***) measures the relative (to reference proteins and each other) amounts of SUUR across fractions. To measure the absolute levels of SUUR, a semi-quantitative approach was used by building a titration curve from SWATH acquisitions of known amounts of recombinant SUUR (***Figure 1-figure supplement 2A&B***). We estimated the amount of SUUR in the nuclear extract (∼140 pg in 25 µg total protein, ***Figure 1-figure supplement 2B***) and in individual fractions from all chromatographic steps (***Figure 1-figure supplement 2C***).

Although in five FPLC steps we achieved >3,000-fold purification of SUUR, it remained only ∼2% pure (***Figure 1-figure supplement 2D***). A progressive loss of material precludes further purification (300 ng of SUUR in 16 µg total protein). Thus, the SUMM4 complex would be nearly impossible to purify to homogeneity from a substantial amount of starting material (∼1 kg *Drosophila* embryos, ∼2.5 g protein), suggesting that SUMM4 could not be identified by the classical FPLC approach.

### SUMM4 regulates the function of *gypsy* insulator elements

Both subunits of SUMM4 contribute to the known functions of *gypsy* insulator (***Figure 4A****-**C***).

Although a *SuUR* mutation decreased the insulator activity, the suppression was universally weaker than that by *mod(mdg4)^u1^*. It is possible that SUUR is not absolutely required for the establishment of the insulator. For instance, the loss of SUMM4 may be compensated by the alternative complex of Mod(Mdg4)-67.2 (***Figure 3-figure supplement 1E****-**H***). Furthermore, the *mod(mdg4)^u1^* allele is expected to have an antimorphic function, since it can mis-localize interacting partner proteins, including SUUR itself (***Figure 3B&C***). Interestingly, *SuUR* has been previously characterized as a weak suppressor of variegation of the *white^m4h^*X chromosome inversion allele, which places the *white* gene near PH (Belyaeva et al., 2003). In contrast, *SuUR* mutation enhances variegation in the context of insulated, PH-positioned *white* (***Figure 3C***). Therefore, this phenotype is unrelated to the putative *Su(var)* function of *SuUR* but, rather, is insulator-dependent.

### ATP-dependent motor proteins are required for the establishment of chromatin barrier and chromosome partitioning

Our discovery and analyses of SUMM4 provide a biochemical link between ATP-dependent motor factors and the activity of insulators in regulation of gene expression and chromatin partitioning.

Insulator elements organize the genome into chromatin loops (Gerasimova et al., 1995) that are involved in the formation of topologically associating domains, TADs (Peterson, Samuelson, & Hanlon, 2021; Rowley et al., 2017; Szabo, Bantignies, & Cavalli, 2019). In mammals, CTCF-dependent loop formation requires ATP-driven motor activity of SMC complex cohesin (Davidson et al., 2019). In contrast, CTCF and cohesin are thought to be dispensable for chromatin 3D partitioning in *Drosophila* (Matthews & White, 2019). Instead, the larger, transcriptionally inactive domains (canonical TADs) are interspersed with smaller active compartmental domains, which themselves represent TAD boundaries (Rowley et al., 2017). It has been proposed that in *Drosophila*, domain organization does not rely on architectural proteins but is established by transcription-dependent, A-A compartmental (gene-to-gene) interactions (Rowley et al., 2017). However, *Drosophila* TAD boundaries are enriched for architectural proteins other than CTCF (Van Bortle et al., 2014), and their roles have not been tested in loss-of-function models. Thus, it is possible that in *Drosophila*, instead of CTCF, the 3D partitioning of the genome is facilitated by another group of insulator proteins, such as Su(Hw) and SUMM4 that together associate with class 3 insulators (Schwartz et al., 2012).

Moreover, SUUR may provide the DNA motor function to promote a physical separation of active and inactive loci and help establish chromosome contact domains (***Figure 4D****-**F***). The translocation model is supported by observations of an asymmetric, selective propagation of SUUR from its initial sites of deposition via Su(Hw)-Mod(Mdg4) binding towards inside of IH regions but not outside (***Figure 4-figure supplement 1E***) (Filion et al., 2010), which may be facilitated by physical interactions between SUUR and linker histone H1 enriched in IH (Andreyeva et al., 2017). It has been reported that another *Drosophila* BTB/POZ domain insulator protein CP190 forms a complex with a DEAD-box helicase Rm62 that contributes to the insulator activity (Lei & Corces, 2006). Thus, ATP-dependent motor proteins may represent an obligatory component of the insulator complex machinery.

### SUMM4 mediates known biological functions of SUUR

Our discovery explains previous observations about biological functions of SUUR. For instance, the initial deposition of SUUR and its co-localization with PCNA has been proposed to depend on direct physical interaction with components of the replisome (Kolesnikova et al., 2013). Our model indicates that instead, the apparent co-localization of SUUR with PCNA throughout endo-S phase (***Figure 3-figure supplement 1J***) may be caused by a replication fork retardation at insulator sites. SUUR is deposited in chromosomes as a subunit of SUMM4 complex at thousands of loci by tethering via Mod(Mdg4)-Su(Hw) interactions. As replication forks progress through the genome, they encounter insulator complexes where replication machinery pauses for various periods of time before resolving the obstacle. Thus, the increased co-residence time of PCNA and SUUR manifests cytologically as their partial co-localization. With the progression of endo-S phase, some of the SUMM4 insulator complexes are evicted and thus, the number of SUUR-positive loci is decreased, until eventually, the replication fork encounters nearly completely impenetrable insulators demarcating the UR domain boundaries.

This mechanism is especially plausible given that boundaries of IH loci very frequently encompass multiple, densely clustered Su(Hw) binding sites (*e.g*., ***Figure 4G***, ***Figure 4-figure supplement 1C***). We examined the data from genome-wide proteomic analyses for Su(Hw) and SUUR performed by DamID in Kc167 cells (Filion et al., 2010). Strikingly, Su(Hw) DamID-measured occupancy does not exhibit a discrete pattern expected of a DNA-binding factor. Instead, it appears broadly dispersed, together with SUUR, up to tens of kbp away from mapped Su(Hw) binding sites (***Figure 4-figure supplement 1E***). Interestingly, when Hidden Markov modeling was applied to the DamID data, Su(Hw), Mod(Mdg4)-67.2 and SUUR occupancies were found to strongly correlate genome-wide in a novel chromatin form (“malachite”) that frequently demarcates the boundaries of IH (Khoroshko et al., 2016). These observations strongly corroborate the translocation model for the mechanism of action of SUMM4. According to this model, upon tethering to DNA-bound Su(Hw), SUMM4 traverses the UR region, which helps to separate it in a contact domain. As DNA within the UR region is tracked by SUUR (***Figure 4F***), it is brought into a transient close proximity with both SUMM4 and the associated Su(Hw) protein, which is detected by DamID (or ChIP) as an expanded occupancy pattern.

The deceleration of SUUR-bound replication forks was also invoked as an explanation for the apparent role of SUUR in the establishment of epigenetic marking of IH (Posukh et al., 2015). We propose that global epigenetic modifications observed in the *SuUR* mutant likely do not directly arise from derepression of the replisome as suggested but, rather, result from the coordinate insulator-dependent regulatory functions of SUUR in both the establishment of a chromatin barrier and DNA replication control (*cf* ***Figure 4E&F***).

### Architectural proteins can attenuate replication forks and regulate replication timing

Our work demonstrates for the first time that insulator complexes assembled on chromatin can attenuate the extent of replication in discrete regions of the salivary gland polylploid genome. Despite distinct cell cycle programs in dividing and endoreplicating cells, the biochemical composition of replisomes in both cell types is identical (Zielke et al., 2013). Therefore, similar insulator-driven control mechanisms for DNA replication are likely conserved in mitotically dividing diploid cells. Our data thus implicates insulator/chromatin boundary elements as a critical component of DNA replication control. Our model suggests that delayed replication of repressed chromatin (*e.g*., IH) during very late S phase can be imposed in a simple, two-stroke mechanism (***Figure 4F***). First, it requires that an extended genomic domain is completely devoid of functional origins of replication.

The assembly and licensing of proximal pre-RC complexes can be repressed epigenetically or at the level of DNA sequence. And second, this domain is separated from flanking chromatin by a barrier element associated with an insulator complex, such as SUMM4. This structural organization is capable of preventing or delaying the entry of external forks fired from distal origins.

An important frequent feature of the partially suppressed UR in *mod(mdg4)* animals is its asymmetry (***Figure 4-figure supplement 1C****-**D***), which is consistent with a unidirectional penetration of the UR domain by a replication fork that fires from the nearest external origin (***Figure 4F***). The SUMM4- dependent barrier may be created as a direct physical obstacle to MCM2-7 DNA-unwinding helicase or other enzymatic activities of the replisome. Alternatively, SUMM4 may inhibit the replication machinery indirectly by assembling at the insulator a DNA/chromatin structure that is incompatible with replisome translocation. This putative inhibitory structure may involve epigenetic modifications of chromatin as proposed earlier (Gaszner & Felsenfeld, 2006), linker histone H1 as shown previously (Andreyeva et al., 2017) and may also be dependent on Rif1, a negative DNA replication regulator that acts downstream of SUUR (Munden et al., 2018).

In conclusion, we used a newly developed MERCI approach to identify a stable stoichiometric complex termed SUMM4 that comprises SUUR, a previously known negative effector of replication, and Mod(Mdg4), an insulator protein. SUMM4 subunits cooperate to mediate transcriptional repression and chromatin boundary functions of *gypsy*-like (class 3) insulators (Schwartz et al., 2012) and inhibit DNA replication likely by slowing down replication fork progression through the boundary element. Thus, SUMM4 is required for coordinate regulation of gene expression, chromatin partitioning and DNA replication timing. The insulator-dependent regulation of DNA replication offers a novel mechanism for the establishment of replication timing in addition to the currently accepted paradigm of variable timing of replication origin firing.

## Materials and Methods

### Recombinant proteins

Recombinant proteins were expressed in Sf9 cells using baculovirus system (SUUR, Mod(Mdg4), EGG and WDE), in *E. coli* (ISWI, ModT antigen and LCMS reference proteins) or obtained from EpiCypher Inc. (human BRG1/SMARCA4).

### Sf9 cells

All baculovirus constructs were cloned by PCR with Q5 DNA polymerase (New England Biolabs) and Gibson assembly with NEBuilder HiFi DNA Assembly Cloning kit (New England Biolabs) into pFastBac vector (Thermo Fisher) under control of polyhedrin promoter. All constructs were validated by Sanger sequencing. Baculoviruses were generated according to the protocol by Thermo Fisher. The baculoviruses were isolated by plaque purification, amplified three times, and their titers were measured by plaque assay. FLAG-SUUR construct was cloned from *SuUR-RA* cDNA (LD13959, DGRC). The following open reading frame (ORF) was expressed: M**DYKDDDDK**H-SUUR-PA(1..962)- VEACGTKLVEKY*. To generate ATPase-dead mutant, SUUR-PA(K59) codon was replaced with an alanine codon by PCR and Gibson cloning. Mod(Mdg4)-67.2-V5-His_6_ and Mod(Mdg4)-59.1-V5-His_6_ constructs were cloned from cDNAs *mod(mdg4)-RT* and *mod(mdg4)-RI* synthesized as gBlocks by IDT, Inc. The following ORFs were expressed: Mod(Mdg4)-67.2(1..610)-GIL**EGKPIPNPLLGLDST**GASVEHHHHHH* and Mod(Mdg4)-59.1(1..541)-GIL**EGKPIPNPLLGLDST**GASVEHHHHHH*. EGG-FLAG and EGG (untagged) were cloned by PCR from *egg-RA* cDNA (IP14531). The following ORF was expressed: EGG-PA(1..1262)-**`DYKDDDDK`**`*` and EGG-PA(1..1262)-`*`. FLAG-WDE was cloned by PCR from *wde-RA* cDNA (LD26050). The following ORF was expressed: M**DYKDDDDK**-WDE-PA(2..1420)-`*`. The sequences of FLAG and V5 tags are highlighted in bold typeface.

Cells, 2•10^6^/ml in Sf-900 II SFM medium (Gibco), were infected at multiplicity of infection (MOI) of ∼10 in PETG shaker flasks (Celltreat, Inc.). After infection for 48-72 hours at 27°C, cells were harvested, and recombinant proteins were purified by FLAG or Ni-NTA affinity chromatography (Fyodorov & Kadonaga, 2003). Whereas, typically, amplified baculovirus stocks had titers above 5•10^9^ pfu/ml, FLAG-SUUR viruses reached no more than 2-4•10^8^ pfu/ml, presumably, due to the inhibitory effect of over-expressed protein on viral DNA replication. Accordingly, whereas typical yields of purified recombinant proteins were >100 µg from 1 L Sf9 cell culture, SUUR polypeptides were produced at no more than 2 µg from 1 L culture, which also adversely affected the protein purity (***Figure 1B&2A*** and ***Figure 2-figure supplement 1A&B***).

### E. coli

The expression construct for untagged recombinant *Drosophila* ISWI was prepared from a full- length ISWI cDNA (Ito et al., 1999). Human TXNRD1 sequence was cloned from a cDNA provided by Addgene (#38863), and TXNRD2 was synthesized as a gBlock gene fragment by IDT, Inc. The ORFs were inserted by Gibson cloning in a pET backbone vector in frame with a C-terminal intein-CBD (chitin-binding domain) tag. Protein expression was induced by IPTG in Rosetta 2 cells, and proteins were purified in non-denaturing conditions by chitin affinity chromatography and intein self-cleavage as described (Emelyanov et al., 2014), followed by anion exchange chromatography (Source 15Q) on FPLC. Detailed cloning and purification methods are available upon request. Note that the cloned human thioredoxin reductase ORFs do not express the C-terminal selenocysteines. They were thus presumed catalytically inactive (Arner, Sarioglu, Lottspeich, Holmgren, & Bock, 1999; Cheng & Arner, 2017) and designated hTXNRD1ci and hTXNRD2ci. They were used exclusively as spike-in mass standards in LCMS acquisitions of *Drosophila* proteins.

Polypeptide corresponding to the C-terminal specific region of Mod(Mdg4)-67.2 was cloned in pET24b vector in frame with a C-terminal His_6_ tag. `M`-Mod(Mdg4)-67.2(403..610)-`GILEHHHHHH*` was expressed in Rosetta 2 and purified by Ni-NTA affinity chromatography in non-denaturing conditions. The polypeptide (ModT) was dialyzed into PBS (137 mM NaCl, 3 mM KCl, 8 mM NaH_2_PO_4_, 2 mM KH_2_PO_4_) and used as an antigen for immunizations (see below). All recombinant proteins were examined by SDS-PAGE along with Pierce BSA mass standards (Thermo Fisher), and their concentrations were calculated from infrared scanning of Coomassie-stained gels (Odyssey Fc Imaging System, LI-COR Biosciences).

### Crude cell extracts

#### Nuclear extract from Drosophila embryos

∼1 kg or ∼200 g wild-type (Oregon R) *Drosophila* embryos were collected 0-12 h after egg deposition (AED) from population cages. The embryos were dechorionated, and nuclear extracts were prepared as described (Kamakaka, Tyree, & Kadonaga, 1991). Protein concentration was measured by Pierce BCA assay (Thermo Fisher). The extracts were fractionated by FPLC (***Figure 1B*** and ***Figure 3-figure supplement 1E***) on AKTA PURE system (Cytiva Life Sciences). Aliquots of chromatographic fractions were examined by quantitative shotgun proteomics or western blot analyses as described below. Peak SUUR or Mod(Mdg4) fractions were diluted to appropriate ionic strength (if applicable) and used as a starting material for the next chromatographic step. Details on FPLC column sizes and run parameters are available upon request.

#### E. coli *lysate*

40-ml Rosetta 2 overnight culture was harvested by centrifugation, resuspended in 20 ml HEG (25 mM HEPES, pH 7.6, 0.1 mM EDTA, 10% glycerol) supplemented with 0.1 M KCl, 1 mM DTT and 2 mM CaCl_2_. Cells were disrupted by sonication and centrifuged to remove insoluble material. Nucleic acids were digested with 15 units micrococcal nuclease (Sigma Aldrich) for 20 min at 37°C, and the proteins were precipitated with 2 M ammonium sulfate. The pellet was resuspended in 10 ml HEG + 0.1 M KCl + 1 mM DTT with protease inhibitors (0.5 mM benzamidine, 0.2 mM PMSF) and dialyzed against the same buffer. After centrifugation, the concentration of soluble protein was measured by BCA assay, the *E. coli* lysate was diluted to 1 mg/ml using 100 mM ammonium bicarbonate (ABC) and stored at -80°C.

### Mass-spectroscopy samples

#### Column fractions

For each chromatographic step, 14 to 20 fractions were selected based on the protein fractionation profile according to the UV (A_280_) absorbances measurements. 50-100 µl aliquots of chromatographic fractions, starting material (SM) and column flow-through (FT, if applicable) were saved, and protein concentrations were estimated based on their UV absorbances (1,000 mU A_280_ was considered to be equivalent to 5 mg/ml total protein). Equal volumes of each fraction, SM and FT were used for MS acquisitions, so that no more than 40 µg total protein was processed in each reaction. As a reference, the reactions were supplemented with 1.5 µg each of purified recombinant human thioredoxin reductases 1 and 2 (hTXNRD1ci and hTXNRD2ci, catalytically inactive) expressed in *E. coli*. Dithiotreitol (DTT) was added to the protein samples to 10 mM and NP-40 – to 0.02%. Reaction volumes were brought to 85 µl with 50 mM ammonium bicarbonate (ABC). All reagents, including water, were HPLC/MS grade. The proteins were reduced for 1 h at 37°C and then alkylated with 30 mM iodoacetamide (IAA, 15 µl 200 mM IAA in water) for 45 min at room temperature in the dark. Alkylated proteins were desalted into 50 mM ABC using ZebaSpin columns (40 kDa MWCO) and digested with 1 µg trypsin for 2 h at 37°C. 1 µg more trypsin was added, and the digestion progressed at 37°C overnight. Tryptic peptides were lyophilized for 2 h on SpeedVac with heat and resuspended in 100 µl Sample Buffer: 1% acetonitrile (ACN) and 0.1% formic acid (FA) in water. Equal volumes (23 µl) of samples were used for IDA and SWATH acquisitions (in triplicate) as described below.

#### Recombinant SUUR

To generate the recombinant SUUR reference spectral library (ILR), ∼0.5 µg purified recombinant FLAG-SUUR (both 130 and 65 kDa bands, ***Figure 1A***) was mixed with 1.5 µg each of hTXNRD1ci and hTXNRD2ci and processed for an IDA acquisition as described above, except for 0.5 µg trypsin was used in each cleavage step, and the peptide sample was resuspended in 30 µl Sample Buffer. For SWATH titration of SUUR (***Figure 1-figure supplement 1H&I***), 1 µg recombinant FLAG-SUUR was mixed with 25 µg *E. coli* lysate protein and 1.5 µg each of hTXNRD1ci and hTXNRD2ci. 10-fold serial dilutions down to 10 fg SUUR were also prepared using the mixture of *E. coli* lysate with reference proteins. The samples were processed for SWATH acquisitions in triplicate as described above, 30 µl of sample per injection.

#### In-gel digestion of recombinant proteins for LCMS identification

Recombinant SUUR or SUMM4 purified by FLAG immunoaffinity chromatography was resolved on SDS-PAGE, stained with Coommassie Blue (***Figure 1A&2A***), and up to eight most prominent protein bands were excised. The gel slices were transferred to 1.5-ml Eppendorf tubes, gently crushed with a RotoDounce pestle and destained with 25 mM ABC in 50% methanol and then with 25 mM ABC in 50% ACN (30 min each at room temperature). The proteins were reduced in 50 µl 10 mM DTT for 1 h at 55°C and alkylated with 30 mM IAA for 45 min at room temperature in the dark. The gel fragments were washed with 25 mM ABC in 50% ACN, dehydrated with 100% ACN, dried in a SpeedVac, rehydrated by addition of 50 µl 50 mM ABC and digested with 0.25 µg trypsin overnight at 37°C. The peptides were extracted once with 50 µl 10% FA and once with 100 µl 3% FA in 60% ACN, both extracts were combined, dried in a SpeedVac and resuspended in 50 µl Sample Buffer. 40 µl of each sample was injected for IDA acquisitions as described below.

### Mass-spectroscopy acquisition methods

LC-MS/MS analyses were performed on a TripleTOF 5600+ mass spectrometer (AB SCIEX) coupled with M5 MicroLC system (AB SCIEX/Eksigent) and PAL3 autosampler.

#### Instrument settings

LC separation was performed in a trap-elute configuration, which consists of a trapping column (LUNA C18(2), 100 Å, 5 μm, 20 × 0.3 mm cartridge, Phenomenex) and an analytical column (Kinetex 2.6 µm XB-C18, 100 Å, 50 × 0.3 mm microflow column, Phenomenex). The mobile phase consisted of water with 0.1% FA (phase A) and 100% ACN containing 0.1% FA (phase B). 200 ng to 10 μg total protein was injected for each acquisition. Peptides in Sample Buffer were injected into a 50-µl sample loop, trapped and cleaned on the trapping column with 3% mobile phase B at a flow rate of 25 μl/min for 4 min before being separated on the analytical column with a gradient elution at a flow rate of 5 μl/min. The gradient was set as follows: 0 to 48 min: 3% to 35% phase B, 48 to 54 min: 35% to 80% phase B, 54 to 59 min: 80% phase B, 59 to 60 min: 80% to 3% phase B, and 60 to 65 min at 3% phase B. An equal volume of each sample (23 µl) was injected four times, once for information-dependent acquisition (IDA), immediately followed by data-independent acquisition (DIA/SWATH) in triplicate. Acquisitions of distinct samples were separated by a blank injection to prevent sample carry-over. The mass spectrometer was operated in positive ion mode with EIS voltage at 5,200 V, Source Gas 1 at 30 psi, Source Gas 2 at 20 psi, Curtain Gas at 25 psi and source temperature at 200°C.

#### Information-dependent acquisitions (IDA) and data analyses

IDA was performed to generate reference spectral libraries for SWATH data quantification. The IDA method was set up with a 250-ms TOF-MS scan from 400 to 1250 Da, followed by MS/MS scans in a high sensitivity mode from 100 to 1500 Da of the top 30 precursor ions above 100 cps threshold (100 ms accumulation time, 100 ppm mass tolerance, rolling collision energy and dynamic accumulation) for charge states (*z*) from +2 to +5. IDA files were searched using ProteinPilot (version 5.0.2, ABSciex) with a default setting for tryptic digest and IAA alkylation against a protein sequence database. The *Drosophila* proteome FASTA file (21,970 protein entries, UniProt UP000000803, 3/21/2020) augmented with sequences for common contaminants as well as hTXNRD1 and hTXNRD2 was used as a reference for the search. Up to two missed cleavage sites were allowed. Mass tolerance for precursor and fragment ions was set to 100 ppm. A false discovery rate (FDR) of 5% was used as the cutoff for peptide identification.

#### SWATH acquisitions and data analyses

For SWATH acquisitions (Zhu, Chen, & Subramanian, 2014), one 50-ms TOF-MS scan from 400 to 1250 Da was performed, followed by MS/MS scans in a high sensitivity mode from 100 to 1500 Da (15- ms accumulation time, 100 ppm mass tolerance, +2 to +5 *z*, rolling collision energy) with a variable-width SWATH window (Zhang et al., 2015). DIA data were quantified using PeakView (version 2.2.0.11391, ABSciex) with SWATH Acquisition MicroApp (version 2.0.1.2133, ABSciex) against selected spectral libraries generated in ProteinPilot. Retention times for individual SWATH acquisitions were calibrated using 20 or more peptides for hTXNRD1ci and hTXNRD2ci. The following software settings were utilized: up to 25 peptides per protein, 6 transitions per peptide, 95% peptide confidence threshold, 5% FDR for peptides, XIC extraction window 20 minutes, XIC width 100 ppm. Protein peak areas were exported as Excel files (***Data S2***) and processed as described below.

### MERCI

MERCI is a novel approach for rapid identification of native protein complexes. It combines enrichment for a target subunit of a putative complex by consecutive FPLC steps and quantitative shotgun proteomics of chromatographic fractions. Crude nuclear extract from *Drosophila* embryos was fractionated as in ***Figure 1A***. At every step, 40 µg or less total protein from each of 10-20 fractions (equal volumes) was supplemented with a fixed amount (1.5 µg each) of exogenous reference proteins (human thioredoxin reductases), reduced, alkylated and digested with trypsin (see above). MS1 and MS2 spectra of tryptic peptides were acquired by IDA, and relative SUUR abundance in fractions was measured by data-independent acquisition (DIA/SWATH) in triplicate. SWATH data were quantified using cognate IDA-derived ion libraries. Protein areas for all quantified proteins were normalized to the sum of those for reference proteins. The relative numbers were averaged across triplicates, with standard deviations calculated. The average numbers for all quantified proteins were further normalized by converting them to Z-scores (see ***Data S2*** for an example of calculations). Peak SUUR fractions (one to five) were then subjected to the next FPLC/MERCI step. After five column steps, the ion library from the ultimate FPLC step (IL5) was used to re-quantify SWATH data from all steps. Z-scores for all purification steps were stitched together, and the large array encompassing all data points for every protein was analyzed by Pearson correlation with SUUR (***Data S2***). The most closely correlated purification profiles served as an indication for protein co-purification, potentially, as subunits of a stable complex.

### Biochemical assays with recombinant proteins

#### Oligonucleosome substrates

Oligonucleosomes were reconstituted *in vitro* as described (Lu et al., 2013) from supercoiled plasmid DNA (3.2 kb, pGIE-0), native core histones and H1 prepared from *Drosophila* embryos (Fyodorov & Levenstein, 2002) by gradient salt dialysis in the presence of 0.2 mg/ml nuclease-free bovine serum albumin (BSA, New England Biolabs). Quality of reconstitution was assessed by MNase and chromatosome stop assays (***Figure 2-figure supplement 1D&E***).

#### ATPase assay

40 nM recombinant proteins were incubated in 25 µl reaction buffer containing 20 mM HEPES, pH 7.6, 0.15 M NaCl, 4 mM MgCl_2_, 1 mM ATP, 0.1 mM EDTA, 0.02% (v/v) NP-40 and 0.1 mg/ml nuclease-free BSA for 60 min at 27°C. Some reactions additionally contained 10 nM pGIE-0 plasmid DNA or equivalent amounts of oligonucleosomes ±H1. ATPase assays were performed using ADP-Glo Max kit (Promega). All reactions were performed in triplicate, the results were normalized to the ADP- ATP titration curve according to the kit manual and converted to enzymatic rates (molecules of ATP hydrolyzed per molecule of enzyme per minute). Averages and standard deviations were calculated. Statistical differences were calculated by Mann-Whitney test.

#### EpiDyne^®^-PicoGreen™ nucleosome remodeling assay

EpiDyne^®^-PicoGreen™ is a restriction enzyme accessibility assay modified for increased throughput and sensitivity (***Figure 2-figure supplement 1F***). In brief, a recombinant ATPase over a concentration range (***Figure 2-figure supplement 1G-J***) was mixed with 10 nM EpiDyne biotinylated nucleosome remodeling substrate (EpiCypher), terminally positioned 6-N-66 (219 bp fragment) or centrally positioned 50-N-66 (263 bp) and 1 mM ATP in 20 µL remodeling buffer, 20 mM Tris-HCl, pH 7.5, 50 mM KCl, 3 mM MgCl_2_, 0.01% (v/v) Tween-20, 0.01% (w/v) BSA. The remodeling reactions were incubated at 23°C in 384-well format. At indicated time points, the reactions were quenched, and nucleosome substrates were immobilized on an equal volume of streptavidin-coated magnetic beads (NEB), pre-washed and resuspended in 2x quench buffer, 20 mM Tris-HCl, pH 7.5, 600 mM KCl, 0.01% (v/v) Tween-20 and 0.01% (w/v) BSA. Beads were successively washed by collection on a magnet (three times with wash buffer, 20 mM Tris-HCl, pH 7.5, 300 mM KCl, 0.01% (v/v) Tween-20) and buffer replacement (once with RE buffer, 20 mM Tris-HCl, pH 7.5, 50 mM KCl, 3 mM MgCl_2_, 0.01% (v/v) Tween-20). Beads were resuspended in 20 µl restriction enzyme mix, 50 units/ml Dpn II (NEB) in RE buffer, and incubated at 23°C for 30 min, collected on a magnet, and supernatants from all wells were transferred to a new plate. They were mixed with an equal volume of Quant-iT™ PicoGreen™ dsDNA reagent (ThermoFisher, Component A) and 1 unit/ml thermolabile proteinase K (NEB) in TE and incubated at 23°C for 1 hr. Fluorescence intensity was detected on an Envision microplate reader with excitation at 480 nm and emission at 531 nm, and data expressed as relative fluorescence units (RFU) through the EnVision Workstation (version 1.13.3009.1409).

### *Drosophila* population culture, mutant stocks and genetics

Wild-type (Oregon R) flies were maintained in population cages on agar-grape juice and yeast paste plates at 26°C, 60% humidity with 12-h dark-light cycle. Mutant flies were reared, and crosses were performed at 26°C on standard cornmeal/molasses medium with dry yeast added to the surface. *SuUR^ES^* was a gift of Igor Zhimulev, and *mod(mdg4)^m9^* was a gift of Yuri Schwartz. All other alleles were obtained from the Bloomington Stock Center, Indiana. Combinations of alleles were produced either by crosses with appropriate balancers and segregation of markers or by female germline meiotic recombination. Intra-chromosomal recombination events were confirmed by PCR of genomic DNA. Details and PCR primer sequences are available upon request.

Fly wings were dissected from ∼5 days old adult males and transferred to the drop of PBS + 0.1% Triton X-100 (PBST). The wings were soaked in 80% glycerol in PBST and photographed using Zeiss AxioVert 200M microscope with EC Plan-Neofluar 2.5X/0.075 lens in bright field and CCD monochrome camera AxioCam MRm. For wing area measurements, images were processed using Fiji/ImageJ2 software package. Statistical differences were calculated by two-tailed t-test, assuming unequal variances. Adult fly eye images were taken on live, CO_2_-anesthetized 2-day-old females on Zeiss stereomicroscope Discovery.V12 using CCD color camera AxioCam MRc.

### Antibodies, immunoblots and immunoprecipitation (IP)

Polyclonal antibody (anti-ModT) was raised in Guinea pigs by Pocono Rabbit Farm & Lab. Rabbit polyclonal antibody to the C-terminus of *Drosophila* XNP/ATRX (anti-XNP) was described previously (Emelyanov et al., 2010). Rabbit and Guinea pig polyclonal antibodies to *Drosophila* SUUR were a gift of Alexey Pindyurin (Nordman et al., 2014) and Igor Zhimulev (Pindyurin et al., 2008). Rabbit polyclonal Mod(Mdg4)-FL antibody to full-length Mod(Mdg4)-67.2 that recognizes all splice forms of Mod(Mdg4) was a gift of Jordan Rowley and Victor Corces. Mouse monoclonal anti-FLAG (M2, Sigma Aldrich), anti-PCNA (PC10, Cell Signaling), anti-β-tubulin and anti-HP1a (E7 and C1A9, Developmental Studies Hybridoma Bank) were obtained commercially.

Western blotting was performed using standard techniques. For FPLC column fraction analyses, 5- 10 µl of starting material and flow-through (if applicable) and 5-15 µl of column fractions were loaded per lane. For expression analyses in salivary glands, 10 salivary glands from L3 larvae of indicated genotype were frozen and thawed, boiled extensively in 40 µl 2x SDS-PAGE loading buffer, centrifuged, and the material equivalent to four salivary glands was loaded per lane. The following dilutions were used: 1:200,000 anti-ModT, 1:1,000 anti-Mod(Mdg4)-FL, 1:1,000 Guinea pig and rabbit anti-SUUR, 1:1,000 anti-HP1a, 1:1,000 anti-β-tubulin and 1:2,000 anti-FLAG. Infrared-labeled secondary antibodies: donkey anti-Guinea pig IRDye 800CW, goat anti-mouse IRDye 800CW, goat anti-rabbit IRDye 800CW, goat anti-rabbit IRDye 680CW and goat anti-mouse IRDye 680RD – were obtained from Li-COR Biosciences and used at 1:10,000. The blots were scanned on Odyssey Fc Imaging System (LI-COR Biosciences).

Immunoprecipitation experiments were performed as described (Emelyanov et al., 2012). 400 µl *Drosophila* embryonic nuclear extracts (∼10 mg total protein) were incubated with 10 µl Guinea pig anti-ModT, 30 µl rabbit anti-SUUR or 20 µl rabbit anti-XNP antibodies for 3 h at 4°C. Immunocomplexes were collected by addition of 25 µl protein A-agarose plus (Thermo Fisher) for 2 h at 4*°*C. After washing four times with 1 ml of buffer HEG (25 mM HEPES, pH 7.6, 0.1 mM EDTA, 10% glycerol) *+* 0.15 M NaCl, the immunoprecipitated proteins were eluted with 80 µl 2x SDS-PAGE loading buffer and analyzed by SDS-PAGE and western blot using Guinea pig or rabbit anti-SUUR and anti-Mod(Mdg4) and mouse anti-HP1a antibodies. For Mod(Mdg4) and HP1a, 8 µl of immunoprecipitated material (equivalent to 1 mg nuclear extract proteins) and 5% input (2 µl nuclear extract, 50 µg total protein) were analyzed. For SUUR, 20 µl of immunoprecipitated material (equivalent to 2.5 mg nuclear extract proteins) and 10% input (10 µl nuclear extract, 250 µg total protein) were analyzed.

### Polytene chromosomes and indirect immunofluorescence (IF) analyses

For all cytological experiments, larvae were reared and collected at 18°C. Polytene chromosomes and whole-mount salivary glands were prepared and analyzed as described previously (Andreyeva et al., 2017). Briefly, salivary glands from wandering third instar larvae were dissected in PBS. Glands were transferred into a formaldehyde-based fixative (one ∼15-μl drop of 3% lactic acid, 45% acetic acid, 3.7% formaldehyde on a coverslip) for 2 min, squashed, and frozen in liquid N_2_. The coverslips were removed, and slides were placed in 70% ethanol for 20 min and stored at −20°C. The slides were washed three times for 5 min in PBST. Primary antibodies were incubated overnight at 4°C in PBST + 0.1% BSA and washed three times for 5 min each with PBST. Secondary antibodies were incubated for 2 h at room temperature in PBST + 0.1% BSA and washed three times for 5 min each with PBST.

DNA was stained with 0.1 μg/ml DAPI in PBST for 3 min, and squashes were mounted in Prolong Glass anti-fade mountant (Molecular Probes). Primary and secondary antibodies were used at the following dilutions: Guinea pig anti-ModT, 1:50,000; rabbit anti-SUUR, 1:100; mouse anti-PCNA, 1:1,000; mouse anti-FLAG, 1:100; Alexa Fluor 488 highly cross-absorbed (HCA) goat anti-mouse, Alexa Fluor 568 HCA goat anti-Guinea pig and Alexa Fluor 647 plus HCA goat anti-rabbit (all Thermo

Fisher), all 1:800. Indirect immunofluorescence (IF) images were obtained with Zeiss AxioVERT 200M microscope and AxioCam MRm mono microscopy camera using a 40x/1.3 Plan-Neofluar or 63x/1.40 Plan-Apochromat lenses with oil immersion. Images were acquired using AxioVision software.

For whole-mount IF staining, L3 larvae were reared at 26°C, and salivary glands were dissected in PBS and fixed in 3.7% formaldehyde (Sigma Aldrich) for 20 min at room temperature. The glands were washed in PBS + 0.3% Triton X-100 and permeabilized for 30 min at 37°C in PBS + 1% Triton X-100. Blocking was performed for 30 min at room temperature in PBS+ 0.3% Triton X-100 supplemented with 10% fetal calf serum and 1% BSA. The glands were incubated with primary antibodies diluted in blocking solution for 48 h at 4°C, washed three times with PBS + 0.3% Triton X-100 for 30 min, and incubated with secondary antibodies in blocking solution overnight at 4°C. The stained glands were washed three times with PBS + 0.3% Triton X-100 for 30 min, stained with DAPI (0.1 μg/ml) for 30 min, and mounted in Prolong Gold anti-fade (Invitrogen). IF images were obtained on a Leica SP8 confocal microscope using a 20X/0.75 PLAPO lens and processed using Fiji/ImageJ software.

To quantify the putative colocalization of SUUR and Mod(Mdg4)-67.2 in polytene chromosomes (***Figure 3A***), the image resolution was reduced to 1,388 x 1,040. Pixel intensities (1,443,520) for SUUR and ModT channels were extracted from Bitmap files (ImageJ), normalized to Z-scores and plotted as an X-Y scatter plot (***Figure 3-figure supplement 1C***). For colocalization analyses, the plot regions (Z_ModT_>1 AND Z_SUUR_<3, green) and (Z_ModT_<1 AND Z_SUUR_>3, red) were excluded from consideration (***Figure 3-figure supplement 1D***).

### Next generation sequencing analyses (NGS)

Salivary glands from female wandering third-instar larvae were isolated and flash-frozen in liquid N_2_ until all samples were collected. Genomic DNA for sequencing was prepared from 25 L3 salivary gland pairs or 10 mg embryos (0-6 h AED) using DNeasy Blood and Tissue kit (Qiagen). Each sample was prepared in triplicate. The tissues were soaked in 180 µl buffer ATL + 20 µl proteinase K (15 mg/ml) and lysed for 2-3 h at 55°C. The reactions were cooled to room temperature, supplemented with 4 µl RNase A, ∼40 mg/ml (Sigma Aldrich), and RNA was digested for 10 min. The genomic DNA was fragmented with 0.002 units DNase I (Thermo Fisher) in 100-µl reactions containing 10 mM Tris-HCl, pH 7.5, 10 mM MnCl_2_, 0.1 mM CaCl_2_, 0.1 mg/ml RNase A and 0.2 mg/ml nuclease-free BSA (1x reaction buffer) for 15 min at 37°C. (DNAse I dilutions were prepared using 1x reaction buffer.) Reactions were stopped by adding 5 µl 0.5 M EDTA, and DNase I was inactivated for 20 min at 65°C. The fragmented DNA was purified on QiaQuick columns using PCR purification kit (Qiagen) and eluted in 40 µl 10 mM Tris-HCl, pH 8.0. The size distribution of DNA fragments (200-600 bp, average ∼400 bp) was confirmed and DNA concentration was measured on 2100 BioAnalyzer (Agilent). Libraries were prepared from 20 ng of fragmented genomic DNA with the ThruPLEX DNA-seq kit using SMARTer® DNA Unique Dual Indexes (TakaraBio) and sequenced 150-bp paired-end reads on an NovaSeq 6000 (Novagene).

The sequencing quality of each sample was assessed using FASTQC version 0.11.7 (Andrews, 2010). Raw paired-end reads were trimmed of adapters using BBDuk from the BBTools software version 38.71 using the parameters: `ktrim=r ref=adapters rcomp=t tpe=t tbo=t hdist=1 mink=11` (Bushnell, 2014). Reads were aligned to the BDGP Release 6 of the *Drosophila melanogaster* genome (dm6) (dos Santos et al., 2015) using Bowtie2 version 2.3.4.1 (Langmead & Salzberg, 2012) and parameters `-q --local --very-sensitive-local --no-unal -- no-mixed --no-discordant --phred33 -I 10 -X 700`. Duplicate reads were marked using Picard 2.2.4 (BroadInstitute) and SAM files were converted to BAM format, filtered for quality (`- bq 5`), and removed of duplicates (`-bF 0x400`) using Samtools version 1.9 (Danecek et al., 2021). To examine replicate concordance, a principal component analysis (PCA) was performed using the deepTools package. Replicates clustered indicating high genome-wide similarity within genotypes (*not shown*). For visualization, replicates were merged (`samtools merge`) and coverage was calculated across 50-bp bins and normalized to counts per million (CPM) using deeptools version 3.2.0: `bamCoverage -bs 50 –normalizeUsing CPM` (Ramirez et al., 2016). Each genotype was scaled to the diploid Oregon R embryo signal in 5-kb bins: `bigWigCompare –-operation first -bs 5000`. DamID-chip data for SUUR and Su(Hw) were retrieved from GSE22069 (Filion et al., 2010). ChIP-chip data for Su(Hw) insulator elements were also used (Negre et al., 2010). UR domains were called using a custom R script to identify regions at least 100 kb in length that fell below the average chromosomal read count as described (Andreyeva et al., 2017). Visualization of all data was performed on the UCSC Genome browser using the dm6 release of the *Drosophila* genome (Kent et al., 2002). Each data set was auto-scaled to its own min and maximum and the data were windowed by mean with 16-pixel smoothing applied.

### Quantitative real-time PCR

Genomic DNA samples prior to DNase I fragmentation (see above) were diluted to ∼0.25 ng/µl.

Real-time PCR was performed using 0.5 ng genomic DNA on a ViiA7 thermocycler (Applied Biosystems) with a three-step protocol (95°C 15 sec, 60°C 30 sec, 68°C 60 sec) and iTaq Universal SYBR Green Supermix (Bio-Rad). Primer sequences are provided in ***Table 2***. Each reaction was performed in three technical replicates for each of the three biological samples (N=9). For each amplicon, the average Ct value (`<CT>`) was calculated and normalized to the average Ct value for a random intergenic genomic sequence as a loading control. Further, for each template, the ΔCt was normalized to the average Ct value for embryonic DNA (diploid control). Standard deviation (0_Ct_) for each reaction in triplicate was also calculated. The following ΔΔCt formula was used: `<ΔΔCt> = (<CTTARGET> – <CTINTERGENIC86D>)SG – (<CTTARGET> – <CTINTERGENIC86D>)embryo`. Standard deviations for <ΔΔCt> were calculated as 0_ΔΔCt_ `= square root of` (0^2^_target_ + 0^2^_intergenic86D_`)/2`. ΔΔCt’s were converted to DNA copy numbers as 2^−<ΔΔCt>^. The confidence interval was calculated in the range _between 2_–<ΔΔCt>–α _and 2_–<ΔΔCt>+α. To examine the putative zygotic function(s) of *SuUR*, heterozygous *SuUR^ES^* parents were produced by balancing with *TM6B, Tb* who were crossed *inter se*. L3 salivary glands were dissected from homozygous *SuUR* mutant progeny, and DNA copy numbers were measured by qPCR as described above.

**Table 2.**
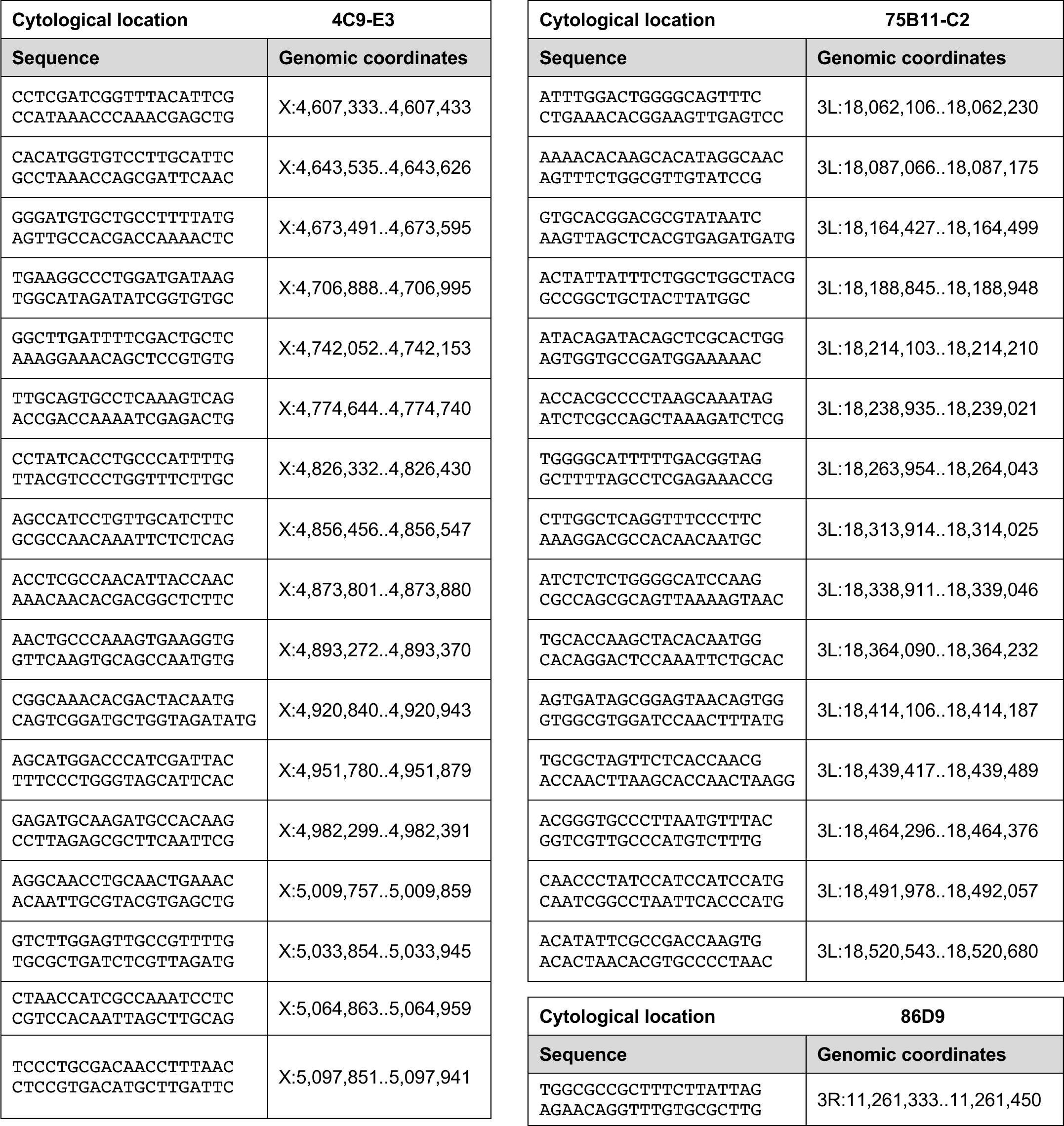
Primer sequences used for qPCR. Genomic coordinates indicate full amplicons, including the length of each primer. Coordinates refer to the BDGP R6/dm3 assembly.

## Acknowledgments

This paper is dedicated to the memory of Jonathan R. Warner who participated in initial discussions that have led to the development of MERCI technology. We thank B. Bartholdy, K. Beirit, B. Birshtein, V. Elagin, M. Gamble, T. Kolesnikova, A. Lusser, A. Pindyurin, C. Schildkraut, Y. Schwartz, S. Sidoli and I. Zhimulev for helpful discussions and critical reading of the manuscript. We thank Y. Schwartz, I. Zhimulev and Bloomington Stock Center for fly stocks and V. Corces, A. Pindyurin, J. Rowley and I. Zhimulev for antibodies. We are grateful to A. Aravin and B. Godneeva for cloning EGG and WDE baculovirus constructs. We thank A. Kumar and N. Baker for help with confocal microscopy, and M. Rogers and J. Secombe for the use of Zeiss Discovery.V12. Confocal images were obtained at the Analytical Imaging Facility (Einstein). We thank P. Schultes for help with maintaining the LCMS instrument.

## Additional Information

### Funding

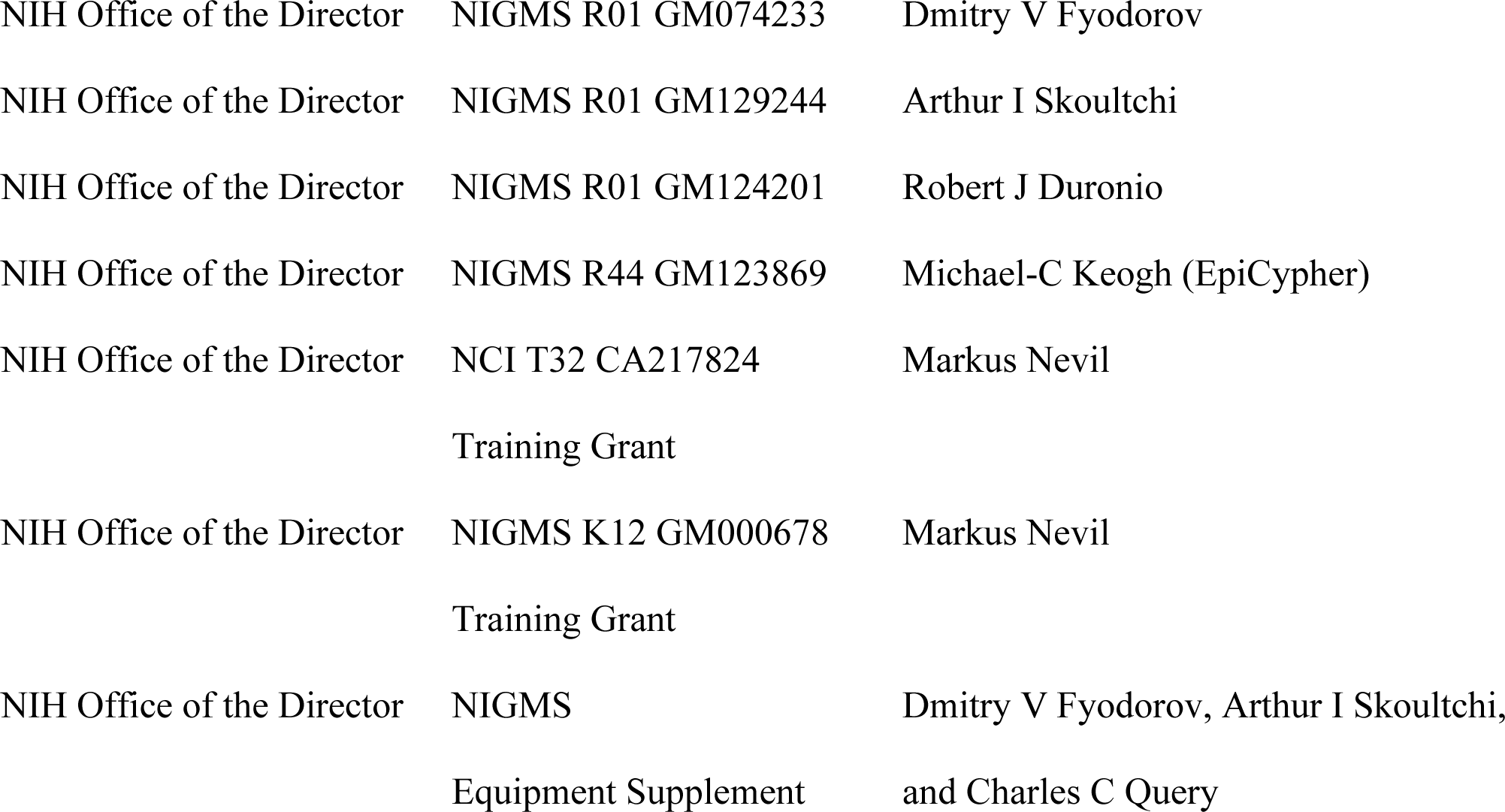

The funders had no role in study design, data collection and interpretation, or the decision to submit the work for publication.

### Author contributions

Evgeniya N Andreyeva, Investigation, Methodology, Validation, Visualization, Interpretation, Writing – review & editing; Alexander V Emelyanov, Conceptualization, Investigation, Methodology, Validation, Visualization, Interpretation, Writing – review & editing; Markus Nevil, Funding Acquisition, Methodology, Validation, Visualization, Interpretation, Writing – review & editing; Lu Sun, Investigation, Methodology, Validation, Visualization, Interpretation, Writing – review & editing; Elena Vershilova, Investigation, Methodology; Christina A Hill, Investigation, Methodology; Michael-C Keogh, Funding Acquisition, Methodology, Project Administration, Resources, Supervision, Validation, Visualization, Interpretation, Writing – review & editing; Robert J Duronio, Funding Acquisition, Methodology, Project Administration, Resources, Supervision, Interpretation, Writing – review & editing; Arthur I Skoultchi, Funding Acquisition, Methodology, Project Administration, Resources, Supervision, Interpretation, Writing – review & editing; Dmitry V. Fyodorov, Conceptualization, Funding Acquisition, Investigation, Methodology, Project Administration, Resources, Supervision, Validation, Visualization, Interpretation, Writing – original draft, Writing – review & editing

### Additional Files

Supplementary Files

Data file S1

Data file S2

### Data Availability

NGS data has been submitted to Gene Expression Omnibus (GEO, accession number GSE189421). *Reviewers may access the data via:* https://www.ncbi.nlm.nih.gov/geo/query/acc.cgi?acc=GSE189421 *using the token upirgumcxngljul*

---

**Figure 1—figure supplement 1.**
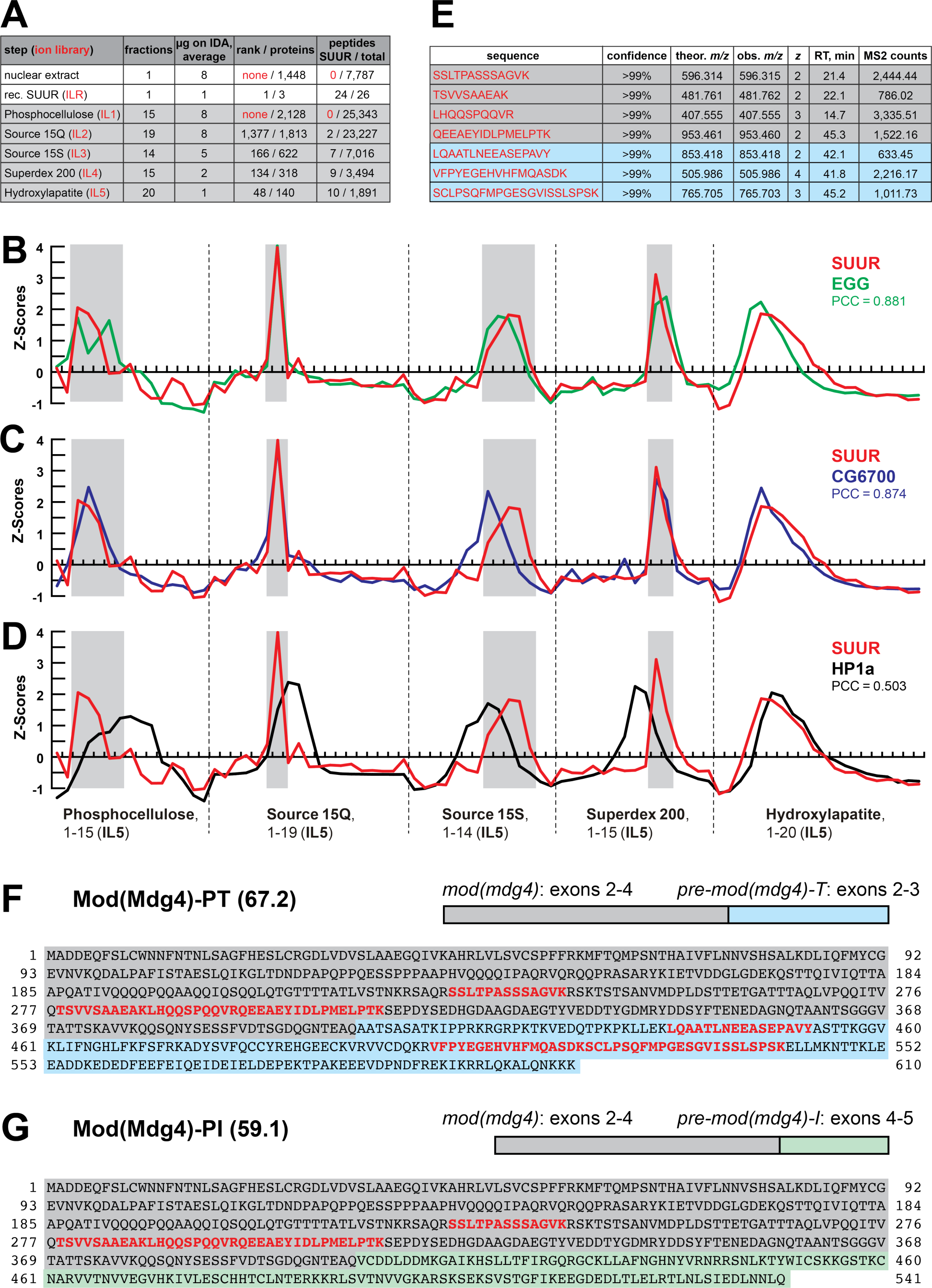
Identification of the SUMM4 complex by MERCI. (A) Representation of SUUR in ion libraries ILR and IL1-5 (Figure 1B, ***Table 1***). Total number of identified proteins and the confidence rank of SUUR among them as well as the total number of detected peptides (95% confidence) and the number of SUUR-specific peptides are shown. (B-D) SWATH quantitation profiles of SUUR (red), EGG (green), CG6700 (blue) and HP1a (black) fractionation across five FPLC steps as in Figure 1D. Pearson coefficients (PCC) are shown (Figure 1E). (E) Mod(Mdg4)-specific peptides from ion library IL5 (***Data S1***). Gray shading, peptides specific to the common part (coding exons 2-4) of Mod(Mdg4); cyan shading, peptides specific to polypeptide Mod(Mdg4)-67.2 encoded by *pre-mod(mdg4)-T*, exons 2-3. (F) Mod(Mdg4)-67.2 polypeptide sequence. The common part is shaded in gray, splice form-specific part is shaded in cyan. Peptides from ion library IL5 (as in E) are highlighted in bold red. (G) Mod(Mdg4)-59.1 polypeptide sequence. The common part is shaded in gray, splice form-specific part is shaded in green.

**Figure 1—figure supplement 2.**
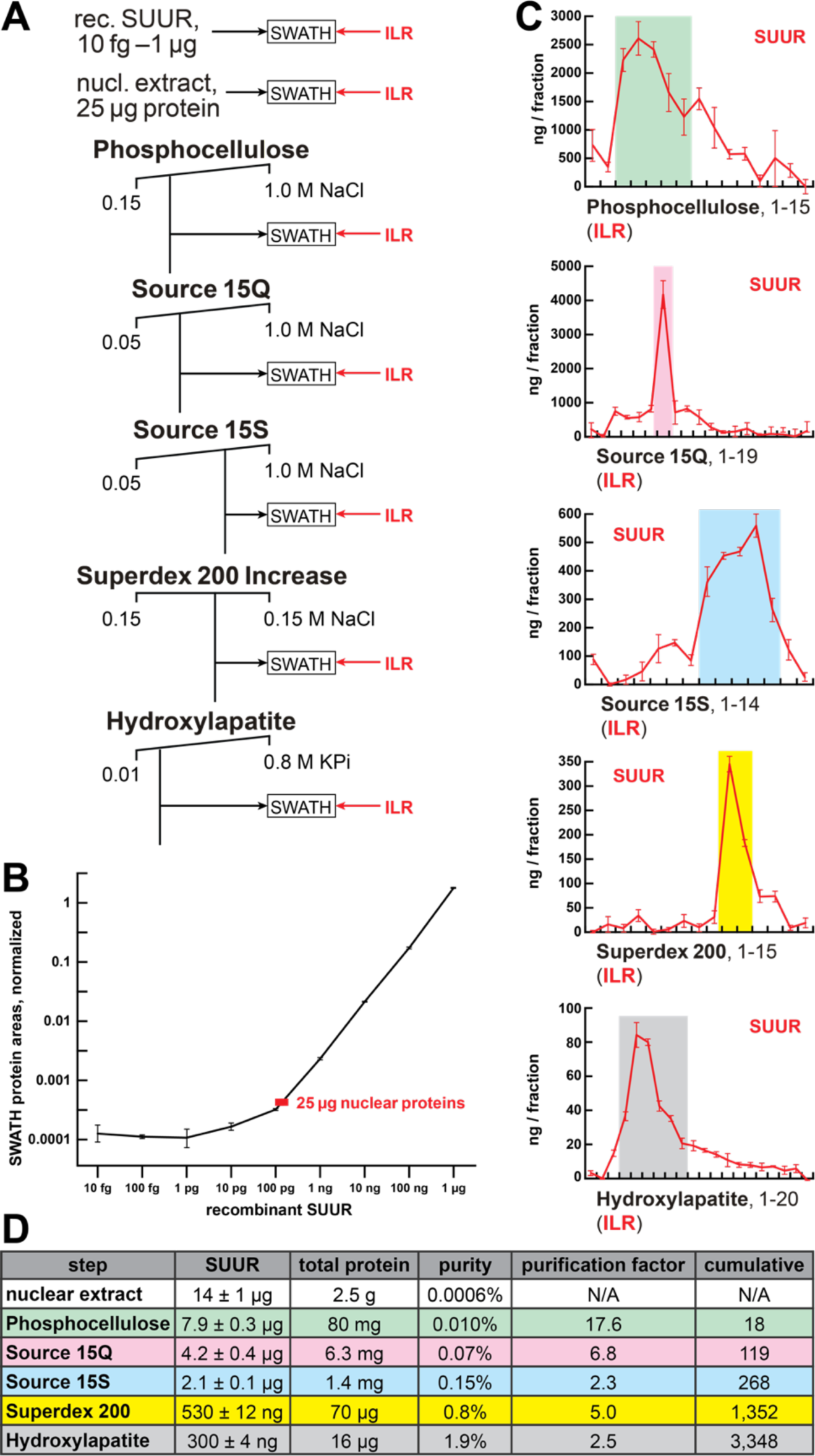
Quantification of SUUR in chromatographic fractions. (A) Schematic of SWATH quantification of recombinant SUUR, nuclear extract (starting material) and FPLC fractions for SUUR using ion library ILR. (B) SUUR titration curve obtained by SWATH quantitation of 10 fg – 1 µg recombinant FLAG-SUUR in the presence of 25 µg *E. coli* lysate; both axes are logarithmic (log_10_). Red rectangle, SUUR quantification in 25 µg nuclear extract; error bars, standard deviations (*N*=3). (C) SWATH quantitation profiles of SUUR fractionation across individual FPLC steps. Ion library ILR was used for SWATH quantification, and relative amounts were converted to estimated ng SUUR per fraction. Error bars, standard deviations (*N*=3); colored boxes, peak fractions of SUUR. (D) SUUR purification by FPLC. Total protein was measured by BCA assay, and SUUR was measured as in (C). Relative purity, purification factor in each step and cumulative purification factor are shown.

**Figure 2—figure supplement 1.**
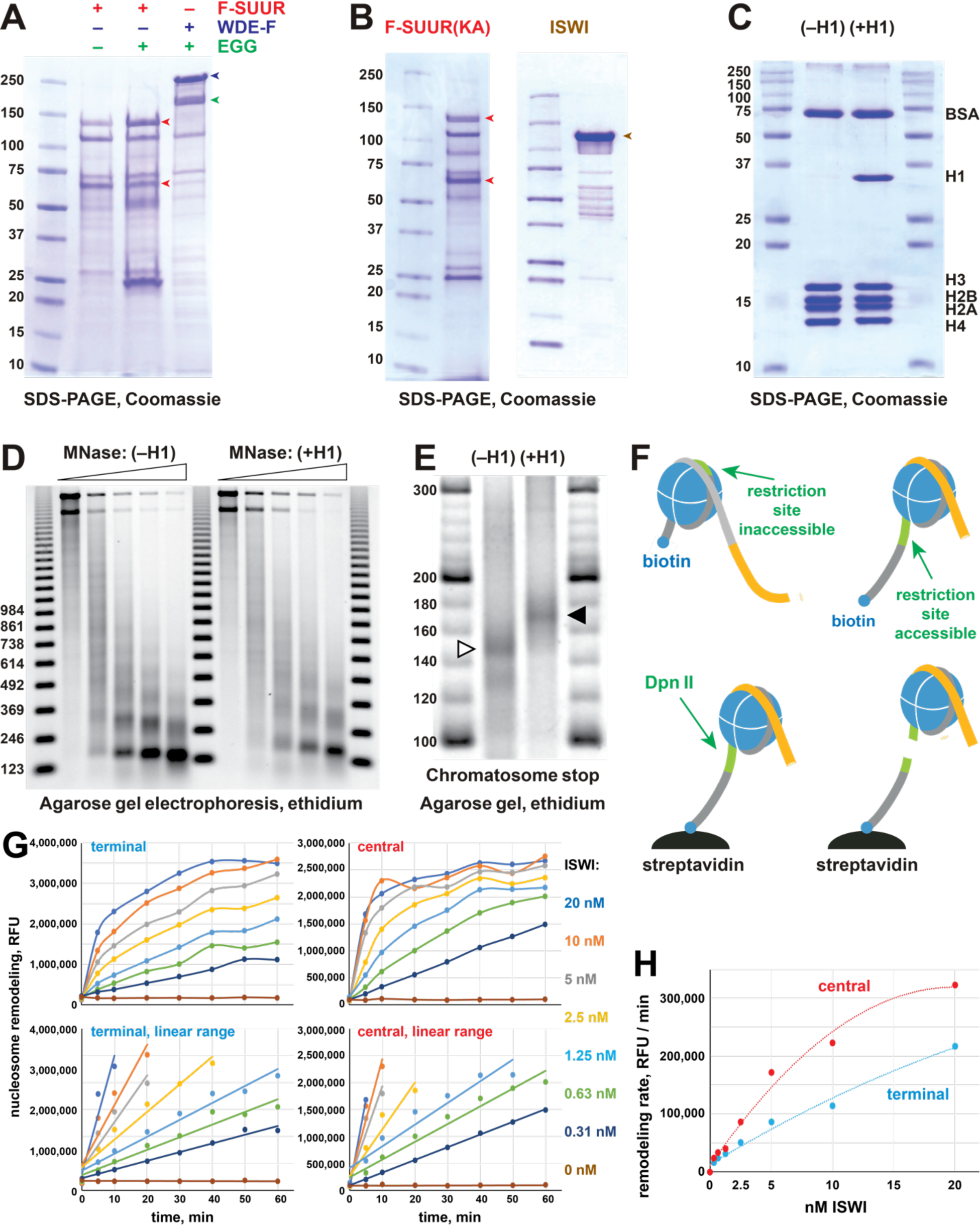
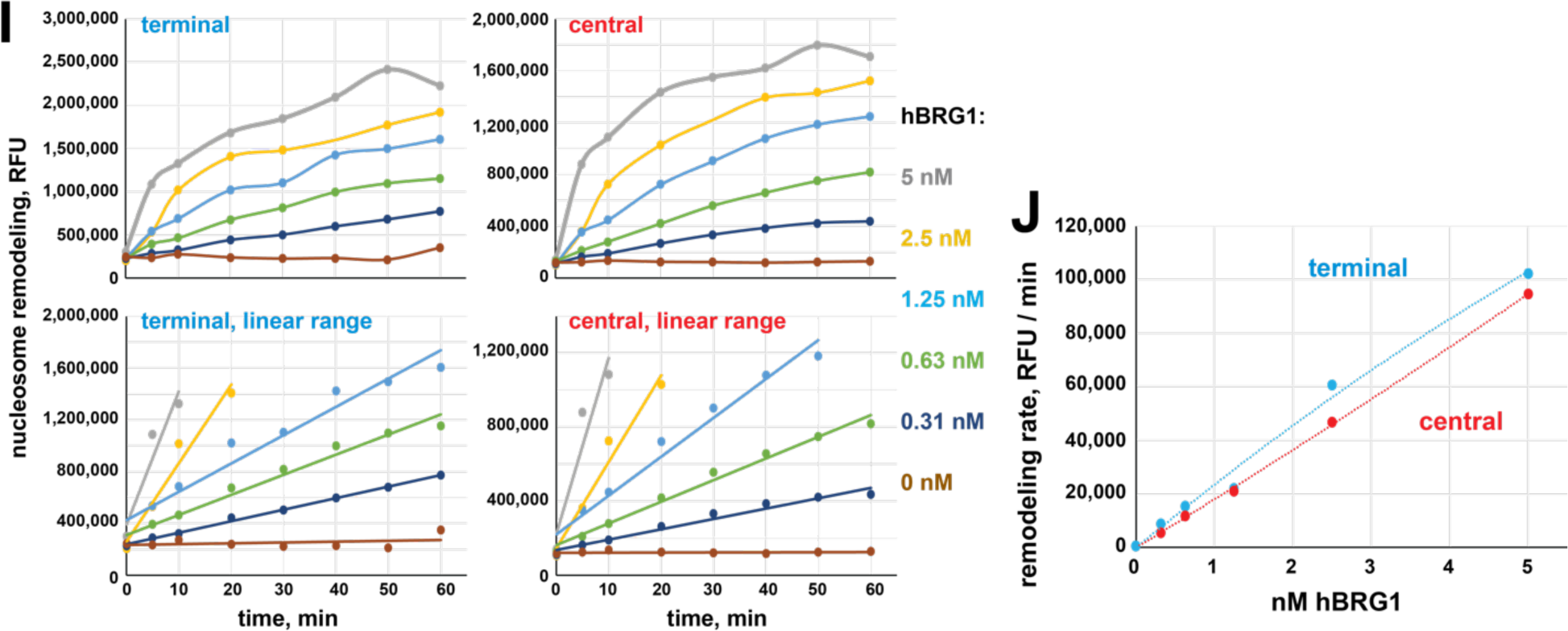
Recombinant proteins, substrates and biochemical assays. (A) Physical interactions of recombinant EGG, SUUR and WDE. Untagged EGG (green arrowhead) was co-expressed with FLAG-SUUR (red arrowheads, p130 and p65) or WDE-FLAG (purple arrowhead) in Sf9 cells and purified by FLAG affinity chromatography. EGG forms a specific complex with WDE but not SUUR. (B) Recombinant FLAG-SUUR(K59A) expressed in Sf9 cells and ISWI expressed in *E. coli*. See legend to ***Figure 1B***. (C) Protein composition of *in vitro* reconstituted chromatin. Oligonucleosomes prepared from plasmid DNA and core histones with (+H1) or without H1 (–H1) were analyzed by SDS-PAGE and Coomassie staining. Positions of BSA, H1 and core histone bands are indicated on the right; molecular mass markers (kDa) are shown on the left. (D) Micrococcal nuclease (MNase) analysis of reconstituted chromatin. Partial digestion with five different dilutions of MNase was performed on H1-free (–H1) and H1-containing (+H1) oligonucleosomes. Deproteinated DNA fragments were analyzed by agarose gel electrophoresis and stained with ethidium. Note the increased nucleosome repeat length in (+H1) lanes consistent with H1 incorporation. Triangles at the top indicate increasing MNase concentrations; 123 bp ladder was used as a molecular mass marker. (E) Chromatosome stop assay. Oligonucleosomes assembled with or without H1 were subjected to partial MNase digestion, and DNA was analyzed by agarose gel electrophoresis and ethidium bromide staining. Positions of the core particle and chromatosome DNA are indicated by arrowheads. DNA fragment sizes in the 20-bp DNA ladder marker are shown. (F) EpiCypher^®^ EpiDyne^®^-PicoGreen™ assay design. EpiDyne nucleosomes encompass a restriction site shielded by the initial nucleosome position but exposed for Dpn II cleavage upon remodeling (sliding or displacement). Biotinylated substrates are immobilized on streptavidin magnetic beads. Digest by Dpn II releases the substrates from beads, and supernatant is quantified by PicoGreen™ (dsDNA detection reagent) fluorescence. (G) Titration of *Drosophila* ISWI remodeling activity using terminally (6-N-66) or centrally (50-N-66) positioned mononucleosomes. Early reaction time points were separately plotted to indicate linear ranges. RFU, relative fluorescence units. (H) Early remodeling rates for ISWI were calculated by linear regression analyses of data in respective linear ranges. ISWI exhibits a stronger remodeling activity with a centrally positioned nucleosome substrate. (I) Titration of human BRG1 remodeling activity. Data are presented as in (G). (J) Early remodeling rates for BRG1 were calculated and plotted as in (H). BRG1 does not exhibit a bias towards remodeling centrally or terminally positioned nucleosomes.

**Figure 3—figure supplement 1.**
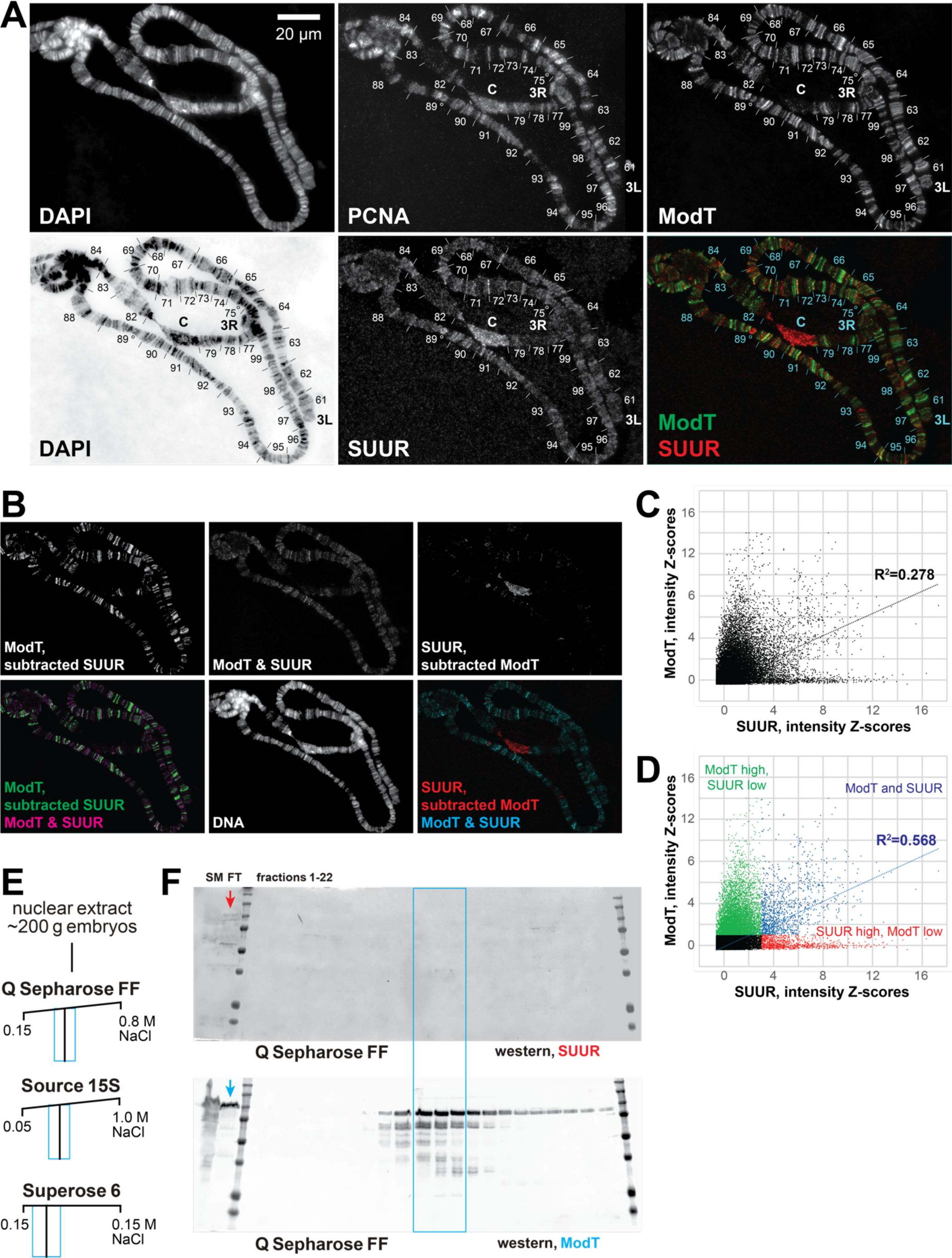
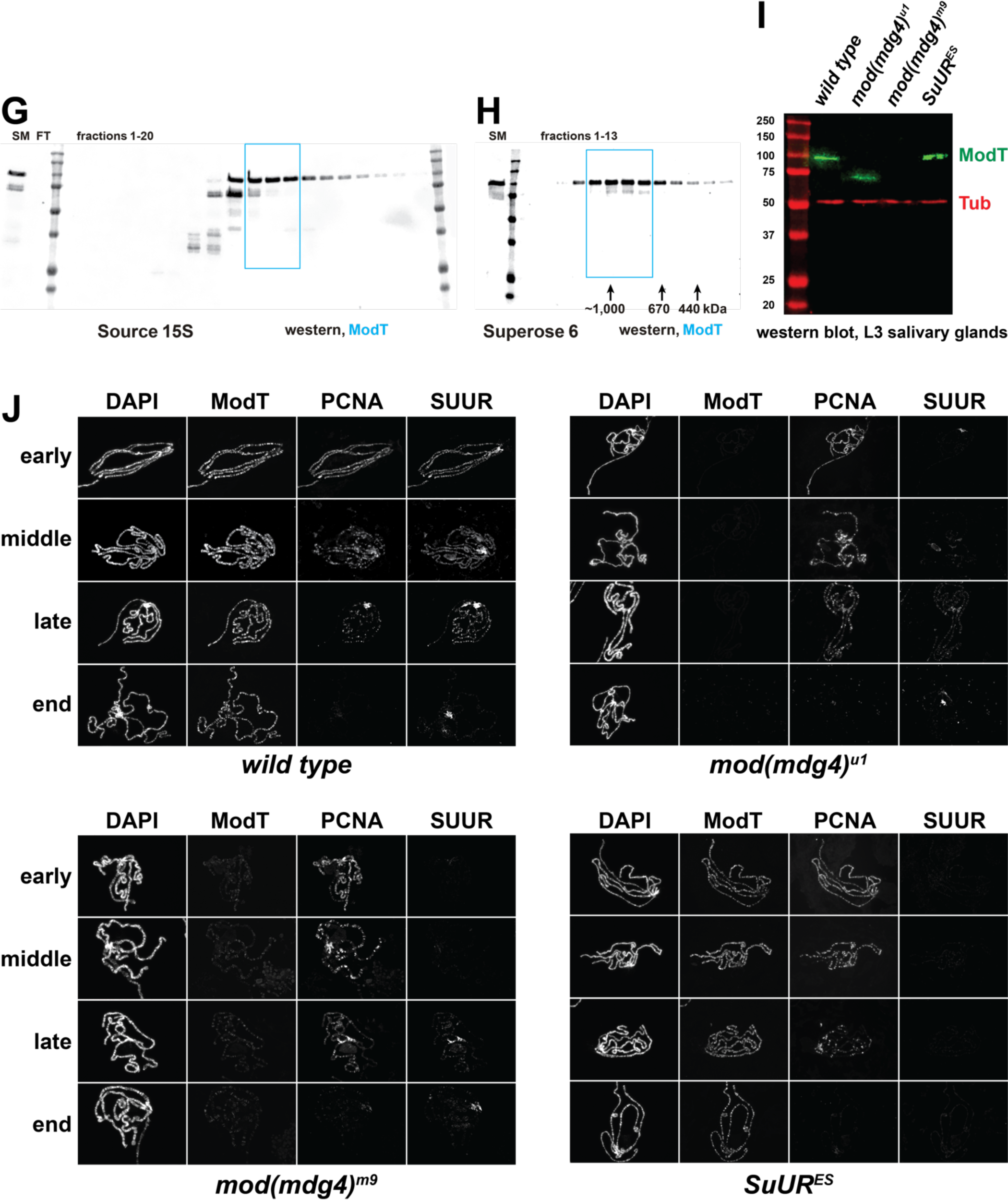
Spatiotemporal distribution of SUUR and Mod(Mdg4) *in vivo* and alternative complex(es) of Mod(Mdg4). (A) Colocalization of SUUR and Mod(Mdg4)-67.2 in *wild-type* polytene chromosomes. See legend to ***Figure 3A***. 3L and 3R telomeres are marked; approximate boundaries of cytological regions are shown according to (Lefevre, 1976); positions of IH regions 75C and 89E that are underreplicated and responsive to *SuUR* mutation are marked by circles. (B) The patterns of colocalization and independent loading of SUUR and Mod(Mdg4)-67.2 in *wild-type* polytene chromosomes. Subtracted and overlapping images were produced in ImageJ (*Supplemental Methods*). Green, enriched Mod(Mdg4)-67.2 and low SUUR; red, enriched SUUR and low Mod(Mdg4)-67.2; magenta or cyan, overlapping enriched Mod(Mdg4)-67.2 and SUUR. (C) Quantification of the overlap between SUUR and Mod(Mdg4)-67.2 in *wild-type* polytene chromosomes (***Figure 3A***). Individual pixel intensities of anti-SUUR and anti-ModT IF signals are normalized to Z-scores and plotted on *x*- and *y*- axes, respectively (*Supplemental Methods*); they exhibit a weak positive correlation (R^2^>0.2). (D) Visually, the 2D plot (C) is split in four separate areas demarcated by Z_ModT_= 1 and Z_SUUR_= 3. When pixels representing ModT-only and SUUR-only areas (green and red, respectively) are removed, the remaining pixels that are simultaneously enriched for Mod(Mdg4)-67.2 and SUUR (blue) exhibit a strong positive correlation (R^2^>0.5). (E) Schematic of partial FPLC purification of an alternative complex of Mod(Mdg4)-67.2. Cyan boxes, fraction ranges used for the next chromatographic step. (F) Western blot analyses of Q Sepharose FF fractions with SUUR and ModT antibodies. SUUR and ∼25% total Mod(Mdg4)-67.2 present in the starting material (SM) fractionate in the flow-through (FT, arrows), whereas Mod(Mdg4)-67.2 also fractionates as an additional, SUUR-free peak (cyan box). Molecular mass markers are as in ***Figure 1G***. (G) Western blot analysis of Source 15S fractions with the ModT antibody. (H) Western blot analyses of Superose 6 fractions with the ModT antibody. Black arrows, expected peaks of globular proteins with indicated molecular masses in kDa. (I) Western blot analyses of lysates of whole salivary glands. L3 salivary glands from homozygous animals of indicated genotypes were probed with ModT (green) and β-tubulin antibodies (red, loading control). Mass marker sizes (kDa) are shown on the left. (J) Spatiotemporal distribution of SUUR in polytene chromosomes. See legend to ***Figure 3B***. Although SUUR is not properly loaded into *mod(mdg4)* chromosomes during early endo-S phase (as in wild type), its deposition partially recovers during late endo-S.

**Figure 4—figure supplement 1.**
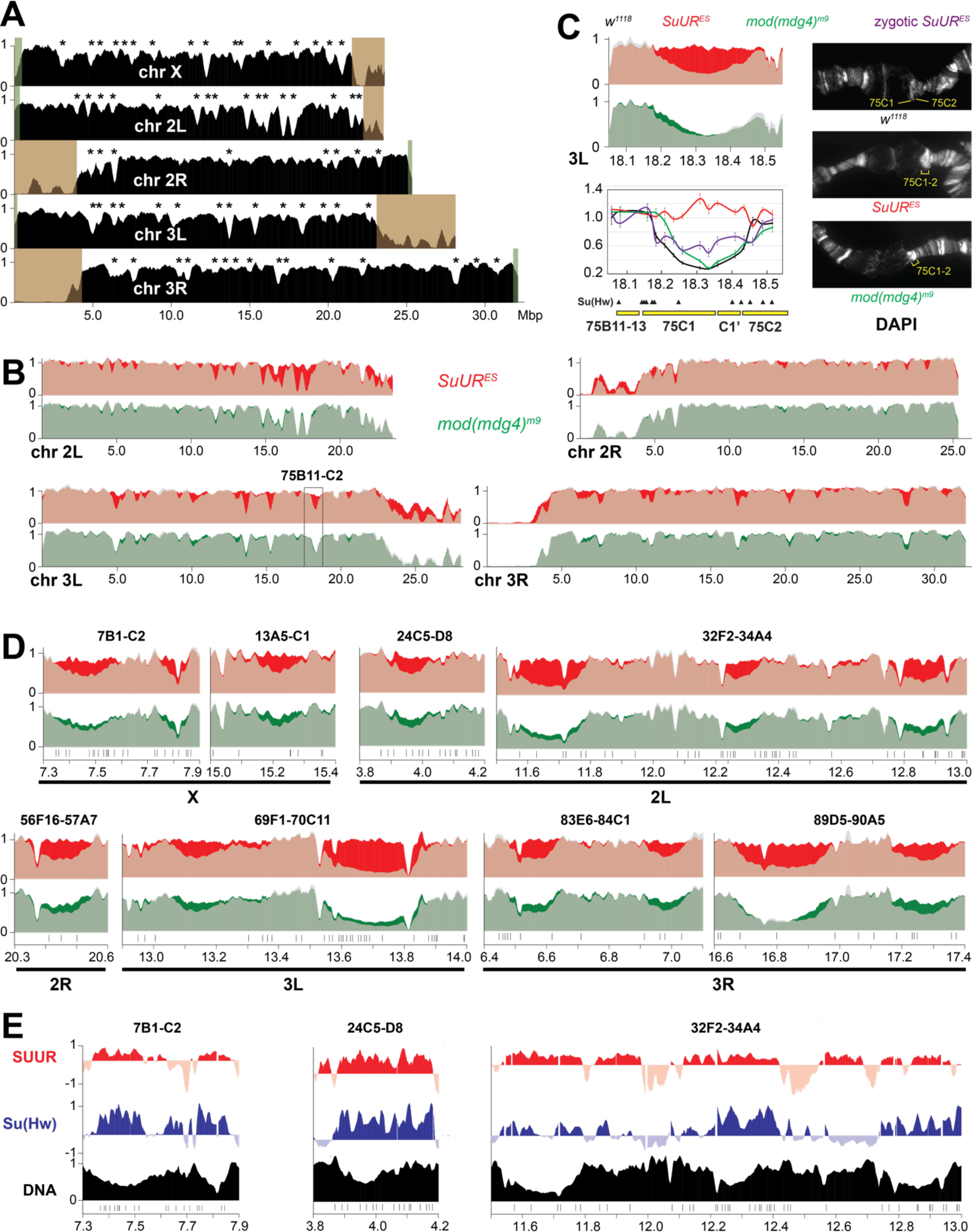
Biological functions of SUMM4 in regulation of under-replication. (A) Genome-wide analyses of DNA copy numbers in *Drosophila* salivary gland cells (*w^1118^* control). Chromosome arms are indicated in white. Brown- and green-shades boxes, mapped pericentric and telomeric heterochromatin regions (Hoskins et al., 2015), respectively. Asterisks, positions of UR domains (Table 1). See legend to ***Figure 4G*** for other designations. (B) Genome-wide analyses of DNA copy numbers in *Drosophila* salivary gland cells in chromosome arms 2L, 2R, 3L and 3R. The data were obtained and presented as for the X chromosome (***Figure 4G***). Black box, 75B11-C2 cytological region. (C) Close-up view of DNA copy numbers by high-throughput sequencing and by qPCR for region 75B11- C2 (data are presented as in ***Figure 4H***) and DAPI-stained polytene chromosome segments around cytological regions 75B-75C. (D) Close-up view of DNA copy numbers by high-throughput sequencing for additional genomic regions. Approximate cytogenetic locations are indicated at the top of each panel. Short vertical bars at the bottom, positions of mapped Su(Hw) binding sites (Negre et al., 2010). See (C) for other designations. (E) Sample plots of DamID profiles for SUUR (red) and Su(Hw) (purple), log_2_ enrichment over Dam-only control (Filion et al., 2010). Positive values are plotted in dark colors and negative values in light colors for contrast. DNA copy numbers in salivary gland cells (black) indicate UR IH domains. Vertical bars, Su(Hw) binding sites (Negre et al., 2010).

## Notes

### Competing Interest Statement

Competing interests
LS and MCK are employed by Epicypher, Inc., a commercial developer and supplier of the EpiDyne nucleosomes and associated remodeling assay platforms used in this study. The remaining authors declare no competing interests.

### Summary of Updates

Fig. 4H was revised.

